# The Arabidopsis *WRR4A* and *WRR4B* paralogous NLR proteins both confer recognition of multiple *Albugo candida* effectors

**DOI:** 10.1101/2021.03.29.436918

**Authors:** Amey Redkar, Volkan Cevik, Kate Bailey, Oliver J. Furzer, Sebastian Fairhead, M. Hossein Borhan, Eric B. Holub, Jonathan D.G. Jones

**Author notes:** Corresponding author: Jonathan D.G. Jones. These authors contributed equally (Amey Redkar, Volkan Cevik).

## Abstract

The oomycete *Albugo candida* causes white blister rust, an important disease of Brassica crops. Distinct races of *A. candida* are defined by their specificity for infecting different host species.

The *White Rust Resistance 4* (*WRR4*) locus in Col-0 accession of *Arabidopsis thaliana* contains three genes that encode TIR-NLR resistance proteins. The Col-0 alleles of *WRR4A* and *WRR4B* confer resistance to at least four *A. candida* races (2, 7 and 9 from *B. juncea, B. rapa* and *B. oleracea*, respectively, and Race 4 from *Capsella bursa-pastoris*). Resistance mediated by both paralogs can be overcome by Col-0-virulent isolates of Race 4.

After comparing repertoires of candidate effectors in resisted and resistance-breaking strains, we used transient co-expression in tobacco or *Arabidopsis* to identify effectors recognized by *WRR4A* and *WRR4B*. A library of CCG effectors from four *A. candida* races was screened for *WRR4A-* or *WRR4B-* dependent elicitation of hypersensitive response (HR). These CCG genes were validated for WRR-dependent HR by bombardment assays in wild type Col-0, *wrr4A* or *wrr4B* mutants.

Our analysis revealed eight *WRR4A*-recognized CCGs and four *WRR4B-*recognized CCGs. Remarkably, the N-terminal region of 100 amino acids after the secretion signal is sufficient for *WRR4A* recognition of these eight recognized effectors. This multiple recognition capacity potentially explains the broad-spectrum resistance to many *A. candida* races conferred by *WRR4* paralogs.

## Introduction

Co-evolution of hosts and their parasites remains a fascinating biological problem. Most plant pathogens fail to colonize most potential hosts, in part because of defense responses initiated upon pathogen perception by cell surface or intracellular immune receptors (Jones & Dangl, 2006). Successful pathogens evade host recognition and suppress defense via effectors (Toruño *et al*., 2016). In co-evolving systems, genetic variation in resistance and susceptibility to particular pathogen races is usually specified by loci encoding Nucleotide Binding, Leucine-rich Repeat (NLR) intracellular immune receptors (Tomborski & Krasileva, 2020). Plant NLRs are under diversifying selection compared to the rest of the genome (Monteiro & Nishimura, 2018). NLR genes are classified as TIR-NLR (TNL) with N-terminal Toll/Interleukin-1/Resistance domains, CC-NLR (CNL) with a coiled-coil (CC) domain, or RPW8-NLR (RNL) with Resistance to Powdery mildeW8 (RPW8) domains (Meyers *et al*. 2003; Zhang *et al*. 2016). Variation in NLR repertoires has been investigated by *Resistance* gene enrichment sequencing (RenSeq) (Jupe *et al*., 2013). NLR immune receptor polymorphism and diversity in 64 accessions of *Arabidopsis thaliana* has been defined using RenSeq (Van de Weyer *et al*., 2019).

NLR-mediated effector-triggered immunity (ETI) often leads to a hypersensitive cell death response (HR) upon effector recognition. Recent work using estradiol-inducible *AvrRps4* activation of ETI in the absence of pattern triggered immunity (PTI), revealed mutual potentiation between PTI and ETI to confer resistance (Ngou *et al*., 2021). Effectors can be recognized via direct interaction with an NLR protein, described as the ‘ligand- receptor model’ (Jia *et al*., 2000; Dodds *et al*., 2006). The flax (*Linum usitatissimum*) L6 NLR protein directly recognizes variants of the flax rust fungus (*Melampsora lini*) effector AvrL567 (Ravensdale *et al*., 2012). Alternatively, some NLRs can detect multiple sequence-unrelated effectors indirectly. Such NLRs either “guard” host proteins that are targeted by multiple effectors (guardee model) (Dangl & Jones, 2001), or guard “decoy” proteins that have evolved to mimic the host target (van der Hoorn & Kamoun, 2008). Effector recognition by some NLRs involves a post-LRR (PL) domain. Recent structural investigations in two TNLs, RPP1 and Roq1 revealed a C-terminal jelly-roll and Ig-like domain (C-JID) in mediating effector binding to form a tetrameric resistosome upon activation (Ma *et al*., 2020; Martin *et al*., 2020).

White blister rust in a wide range of crop and wild Brassica species is caused by oomycete pathogens in the genus *Albugo* (Holub *et al*., 1995; Voglmayr & Riethmüller, 2006; Choi *et al*., 2007). The disease symptoms of white pustules resemble sporulation of basidiomycete rust fungi. Dispersal is *via* dehydrated sporangiospores (Heller & Thines, 2009). Jouet *et al*. (2019) verified that different phylogenetic races of *A. candida* exhibit distinct host specificities for infecting different crop and wild species of Brassicaceae. Important examples include Races 2, 7 and 9 from major crop species (*B. juncea, B. rapa* and *B. oleracea*, respectively) and Race 4 from the common weed *Capsella bursa-pastoria*. Genome comparisons revealed ancient genetic exchange and introgression amongst these *A. candida* races (McMullan *et al*., 2015). *A. candida* species have a remarkable capacity to suppress host resistance to other infections (Cooper *et al*., 2008; Belhaj *et al*., 2017; Prince *et al*., 2017) which creates an immune-compromised state in the colonized host enabling growth of non-adapted pathogens.

*White Rust Resistance 4* (*WRR4*) from *A. thaliana* confers resistance to *A. candida* (Borhan *et al*., 2008). The locus contains three paralogs in *A. thaliana* accession Columbia (Col-0) that encode TNLs. The Col-0 allele of *WRR4A* confers resistance to four *A. candida* races in *A. thaliana* (Borhan *et al*., 2008), and also as a transgene to Ac2V in Brassica crops (Borhan *et. al*, 2010; Cevik *et al*., 2019). The Col-0 allele of *WRR4B* also confers resistance to Brassica-infecting *A. candida* races Ac2V and AcBoT (Cevik *et al*., 2019). Although resistance in Col-0 appears to be broad spectrum, *WRR4* resistance-breaking strains (*e*.*g*., AcEx1) have been collected from white rust pustules on floral stems of wild *A. thaliana* or *A. halleri*. These are natural pathotypic variants of *A. candida* Race 4 (Fairhead, 2016; Jouet *et al*., 2019). Resistance to this Race 4 pathotype occurs in *A. thaliana*, such as in the accession Oystese (Oy-0) which also maps to the *WRR4* locus (Fairhead, 2016; Castel *et al*., 2021).

*A. thaliana* accession Wassilewskija (Ws-2) is resistant in leaves to *A. candida* Race 2 and 7. The *WRR4* locus in Ws-2 is disrupted compared to Col-0 by deletions or sequence variation of *WRR4* paralogs. Ws-2 contains two divergent paralogs (Van de Weyer *et al*., 2019) and one of these (a Ws-2 allele of *WRR4B*) confers resistance to isolates of *A. candida* Race 2 (from *B. juncea*). Both Col-0 and Ws-2 alleles of *WRR4B* also confer resistance in transgenic Brassicas (Cevik *et al*., 2019). Conceivably, allelic variation of TNL paralogs at the *WRR4* locus provides multiple genes that could control white rust in major Brassica crops. Stacking multiple *WRR* genes should promote durability. Thus, identifying and understanding function of *A. candida* effectors recognized by alleles of *WRR4A* and *WRR4B* would help choose the most effective transgene combinations to use for *Albugo* control in Brassica crops.

Recognized effectors from the Peronosporales (including species of *Phytophthora, Pythium* and downy mildews) are translocated into host cells (Kamoun, 2006; Haas *et al*., 2009). These effectors typically carry an N-terminal signal peptide for secretion and an RxLR motif that is implicated in host translocation (Whisson *et al*., 2007; Dou *et al*., 2008). However, in the *P. infestans* effector *Avr3a*, the RxLR motif is cleaved during secretion, and how it promotes translocation still remains unclear (Wawra *et al*., 2017). RxLR effectors show enhanced polymorphism and positive selection for diversification towards their C termini (Rehmany *et al*., 2005; Win *et al*., 2007).

Genome analysis of two *Albugo* species (*Albugo laibachii* and *A. candida*) revealed a new class of oomycete effector-like proteins that carry a “CHxC” motif (Kemen *et al*., 2011). *Albugo* lacks RxLR-encoding proteins compared to the Peronosporales. Re-sequencing of an *A. candida* race from *B. juncea* (Ac2V) using long reads revealed massive expansion of effector-like proteins with a CHxC-like motif, which was reclassified as CX_2_CX_5_G and abbreviated to CCG, resembling previously identified CHxC proteins in *A. laibachii* (Kemen *et al*., 2011; Furzer *et al*., 2021). CCG proteins show no homology to other oomycete secreted proteins. Their high sequence divergence resembles that of RxLR effectors (Furzer *et al*., 2021), and supports their investigation as candidates for being the effectors recognized by *White Rust Resistance (WRR*) genes.

In this study, we compared the genomes of resisted and resistance-breaking strains of *A. candida* to define candidate differentially-recognized CCG effectors. We screened a library of CCG secreted proteins, mainly from *A. candida* races 2 and 4, to identify potential effectors recognized by Col-0 alleles of *WRR4A* and *WRR4B. Agrobacterium*- mediated transient co-expression was used in a pre-screen to identify pairwise combinations of effector and *R-*allele that activated an HR. Twelve CCG candidates were identified including eight recognized by Col-*WRR4A* and four by Col-*WRR4B*. Positive candidates were then validated for HR recognition by a bombardment assay in Col-0 wild type and mutants (*wrr4A* or *wrr4B*) of *A. thaliana*. Several of these CCGs are absent or else show expression polymorphism in the Col-0 virulent isolate AcEx1. To further characterize this *WRR4* recognition, we focused on *WRR4A-*recognized CCGs, for which the N-terminal region is sufficient for recognition. Our data reveal a novel capacity for recognition of multiple effectors of *A. candida* by two distinct *WRR4* paralogs.

## MATERIALS AND METHODS

### Plant material and growth conditions

Wild type and mutant *A. thaliana* accessions used in this study included Col-0, Wassilewskija-2 (Ws-2), Col-0_*wrr4a-6* (Borhan *et al*., 2008), Col-0_*wrr4b* (Cevik *et al*., 2019) and the recombinant inbred line (RIL) CW20 that was derived from a cross between Col-5 x Ws-2 and in this RIL, *WRR4* locus is the only known *WRR* locus introgressed from Col-5 (Fairhead, 2016). Seeds were sown directly on compost and were grown at 21°C, with 10 hours of light and 14 hours of dark, at 75% humidity. For *N. tabacum* and *N. benthamiana*, plants were grown on compost at 21°C, with cycles of 16 hours of light and 8 hours of dark, at 55% humidity.

### *A. candida* infection assay

For leaf inoculations, zoospores harvested from previous leaf infections, were suspended in water (∼10^5^ spores/ml) and incubated on ice for 30 min for releasing of the zoospores from sporangia due to cold shock. The spore suspension was then sprayed on plants using a Humbrol spray gun (Hornby Hobbies Ltd, Sandwich, UK) with a volume equal to ∼700 μl/plant and plants were incubated at 4°C in the dark overnight for efficient zoospore germination. Infected plants were kept under 10-hour light (21°C) and 14-hour dark (16°C) cycles. Phenotypes were monitored between 7 to 10 days post inoculation (dpi) and macroscopic symptoms were readily visible during this period.

The sequential infection assay was done as described previously in McMullan *et al*., 2015. We developed *A. candida* race specific PCR primers by comparing the genome sequences for regions that were unique to the tested *A. candida* races AcNc2 and AcEx1. (Table **S1**). Primers were designed to amplify these regions from genomic DNA extracted from each isolate. Primary inoculum was sprayed onto control and test plants. In the case of AcEx1 *WRR4A* mediated defense suppression assays, both *A. thaliana* Ws-2 and CW-20 were inoculated. The inoculated plants were incubated in the dark at 4°C overnight. Same number of plants treated with water served as mock control. At 7 dpi a secondary infection with the avirulent *A. candida* race AcNc2 was performed on 50% of the plants while the remaining 50% were again mock inoculated with water. The co-inoculated plants were returned to the growth cabinet and incubated for a further 8 dpi. Tissue was harvested and washed in sterile water to remove surface adhering spores, and flash frozen in liquid N2. DNA was prepared using a DNeasy Plant Mini Kit (Qiagen, CA, USA) as described in manufacturer’s instructions. PCR was performed using race-specific primers and products were visualized on a 1% agarose gel.

### Gene cloning and plasmid construction

Cloning of genes were carried out using Uracil-Specific Excision Reagent (USER) method (Geu-Flores *et al*., 2007). Genes with 5’ and 3’ regulatory sequences were cloned into LBJJ233-OD vector (containing a FAST-Red selectable marker) pre-linearized with *Pac*I and Nt.*Bbvc*I. For overexpression, plant genes and CCG effectors, which lack introns, were cloned into pre-linearized LBJJ234-OD (containing a FAST-Red selectable marker, CaMV 35S promoter and Ocs terminator) or pICH86988 (containing Kan selectable marker, CaMV 35S promoter and Ocs terminator). Genes were C-terminally tagged either with a His-FLAG (HF tag) or a yellow fluorescent protein (YFP) tag. Briefly, the candidate CCG effector was PCR amplified from one of the *A. candida* races (AcNc2, AcEm2 or Ac2V) for the high throughput screen for *WRR4A-* recognized CCGs and from race Ac2V for *WRR4B-* recognized CCGs. The genomic DNA was used as a template with KAPA HiFi Uracil+ enzyme, following the manufacturer’s protocol. To obtain mutant versions of CCG28^AAs28-130^ -YFP carrying mutations in the CCG motif (CCG exchanged to AAG, CAA, CAG and AAA), site directed mutagenesis was performed with QuikChange Multi Site-Directed Mutagenesis Kit (Stratagene, Santa Clara, USA) following manufacturer’s instructions. A list of primers and vectors are indicated in Table **S1**. *WRR4A* and *WRR4B* under 35S promoter were cloned in pICH86988 from the Col-0 genomic DNA. All the plasmids were transformed in to *Escherichia coli* DH10B electro-competent cells selected with appropriate antibiotics and purified using a Qiaprep Spin Miniprep Kit (Qiagen). Positive clones were transformed in *Agrobacterium tumefaciens* strain GV3101 and used in infiltrations for transient expression experiments.

### Transient expression in *N. tabacum* or *N. benthamiana* leaves and cell death assay

*A. tumefaciens* strains were streaked on selective media and incubated at 28 °C for 24 hours. The streaked inoculum was transferred to liquid LB medium with appropriate antibiotic and incubated at 28 °C for 24 hours in a shaking incubator at 200 rotations/min (rpm). The resulting cultures was centrifuged at 3,000 rpm for 5 min and resuspended in infiltration buffer (10 mM MgCl_2_, 10 mM MES, 150 μM acetosyringone pH 5.6) at OD_600_ of 0.4 (2 ×10^8^ cfu/ml). For co-expression, each bacterial suspension was adjusted to OD_600_ of 0.4. The abaxial surface of 4-weeks old *N. tabacum* or 5 weeks old *N. benthamiana* were infiltrated with 1 ml needleless syringe. Cell death was phenotyped two to four days after infiltration. For the *WRR4B* recognition assay, macroscopic cell death phenotypes were scored according to the HR index modified from (Segretin *et al*., 2014) ranging from 0 (no visible necrosis) to 6 (full necrosis).

### Particle bombardment in *A. thaliana* and luciferase assay

Transient protein expression in *Arabidopsis* leaves was performed by biolistic gene transfer. 1.0 µm Tungsten particles (Bio-Rad) were coated with the plasmids coding for the indicated CCG genes driven under CaMV 35S promoter (Table **S1**). Bombardment was performed using a PDS-1000/He system (Bio-Rad) onto 4-weeks-old *Arabidopsis* leaves. After bombardment the leaves were incubated in small vials with the leaf petiole immersed in water, for 48 hours post bombardment (hpb). The leaves were then frozen in liquid N2 and stored at -80 °C until further processing.

For the luciferase assay a Dual Reporter Luciferase Assay system (Promega) was used. Four transiently bombarded leaf events were pooled together and crushed in lysis buffer. The extract was centrifuged at 12,000 rpm for 10 min at 4 °C. 20 µl of the lysate was then dispersed in 96 well plate in triplicates and analyzed on Varioskan Flash Instrument by injecting 100 µl of luciferase assay reagent II, which includes substrate and reaction buffer. A 10 second read time was used to measure luciferase activity for each well.

### Gene expression measurement by RT-qPCR

For gene expression analysis, RNA was isolated from three biological replicates and used for reverse transcription quantitative PCR (RT-qPCR) after cDNA synthesis. Briefly, RNA was extracted using the RNeasy Plant Mini Kit (Qiagen) with the DNase treatment (Qiagen). Reverse transcription was carried out using the SuperScript IV Reverse Transcriptase (ThermoFisher). RT-qPCR was performed using CFX96 Touch Real-Time PCR (Bio-Rad). Primers for qPCR analysis of different CCGs are enlisted in Table **S1**. Data were analysed using the double ΔΔCT method (Livak & Schmittgen, 2001) by calculating the relative expression of candidate CCG in relation to the *A. candida EF1α* as a house keeping reference gene.

### Protein extraction and Western Blot

Protein was extracted from *Agrobacterium* infiltrated *N. benthamiana* leaves at 72 hpi as previously described (Sarris *et al*., 2015). Briefly, leaves were harvested and ground in liquid N2, and extracted in GTEN buffer (10% glycerol, 100 mM Tris-HCl, pH 7.5, 1 mM EDTA, 150 mM NaCl, 5 mM 1,4-dithiothreitol (DTT), 1× Complete protease inhibitor mixture (Roche) and 0.2% (V/V) Noniodet P-40). 30 μl of the supernatant from the sample extract was used to elute the samples by boiling in loading buffer. For SDS-PAGE, samples were heated for 10 min at 95 °C for denaturation. After electrophoresis, separated proteins were transferred to PVDF (Merck) membranes for immunoblotting. Membranes were blocked for 2 hours in 5% nonfat milk, probed with horseradish peroxidase (HRP)-conjugated antibodies for overnight. Chemiluminescence detection for proteins was carried out by incubating the membrane with developing reagents (SuperSignal West Pico & West Femto), using ImageQuant LAS 4000 (Life Sciences, USA).

### Statistical analysis

Statistical analysis was carried out using the GraphPad Prism 9.0 software (San Diego, USA). The statistical test employed is provided in the figure legends.

## RESULTS

### The *WRR4A*^Col-0^ and *WRR4B*^Col-0/Ws-2^ provide broad spectrum resistance to *A. candida* races

*WRR4A*^Col-0^ confers full green resistance (GR) to multiple *A. candida* races (Cevik *et al*., 2019; Borhan *et al*., 2008) including the Race 2 isolate Ac2V and two highly similar Race 4 isolates AcNc2 and AcEm2 (McMullan *et al*., 2015) (Fig. **1** and Table **S2**). The *Arabidopsis* accession Ws-2 which lacks *WRR4A* is fully susceptible to AcEm2 but Col-0-*wrr4-6* mutant shows necrotic resistance (NR) rather than GR as observed with wild-type (WT) Col-0, due to the presence of additional *WRR* genes in Col-0. This suggests *WRR4A*^*Col-0*^ contributes to full GR to AcEm2 (Fig. **1**). A Race 4 isolate AcEx1 overcomes *WRR4A*^Col-0^ mediated resistance but triggers chlorosis in infected adult Col-0 plants (Fig. **1**). The Col-0*wrr4-6* mutant exhibits full susceptibility to AcEx1 indicating that *WRR4A*^Col-0^ confers partial resistance to AcEx1.

**Figure 1.**
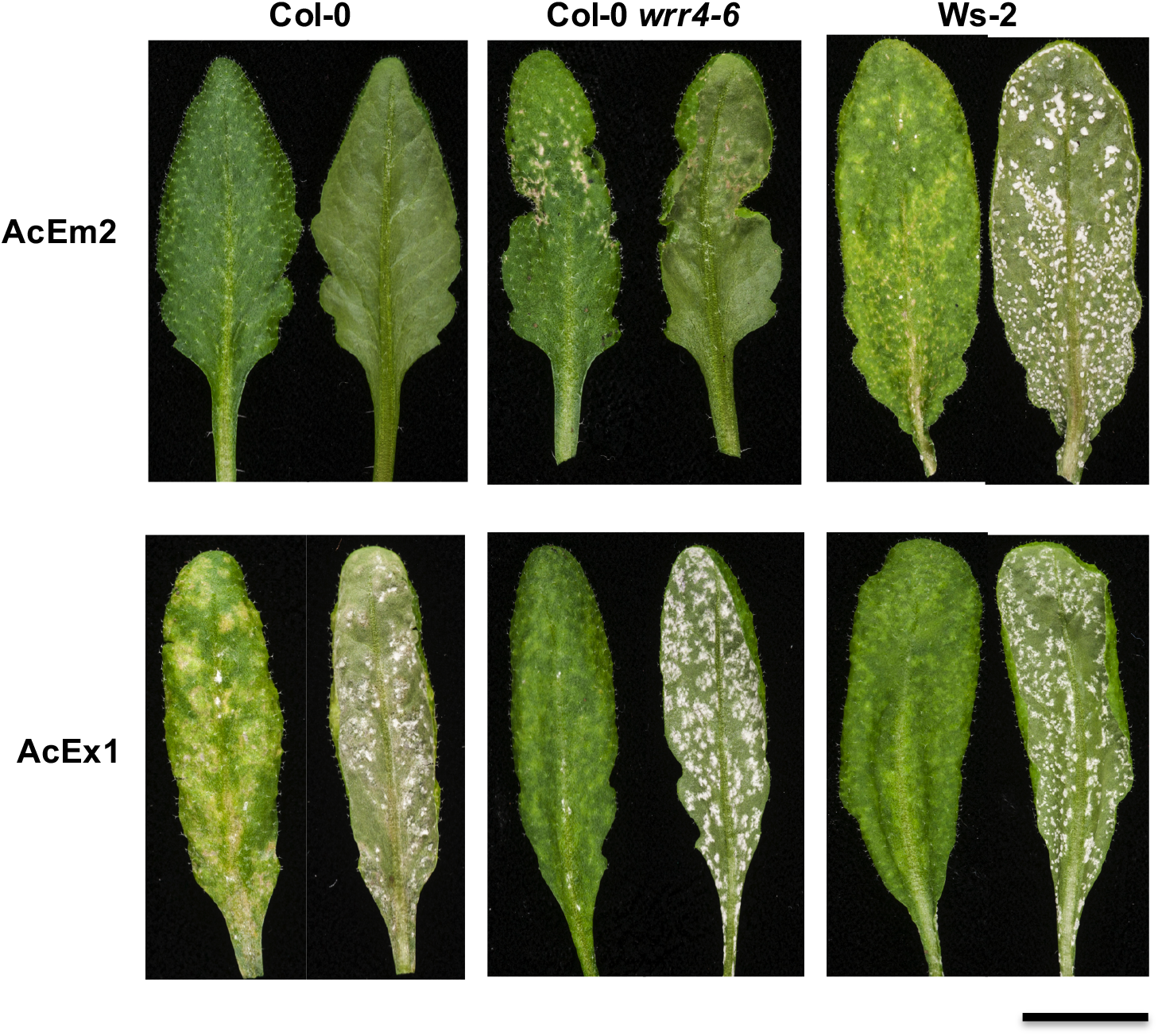
Resistant and susceptible phenotypes of *WRR4A*^Col-0^ *A. thaliana* following inoculation of adult leaves with two different isolates of *A. candida* Race 4. Arabidopsis Col-0 is resistant to Race 4 isolate AcEm2 (collected from *Capsella bursa-pastoris)* and susceptible to Race 4 isolate AcEx1 (collected from *A. halleri*) whereas Ws-2 is susceptible to both isolates. Necrotic resistance observed with the Col-0-*wrr4-6* mutant indicates that *WRR4A* is responsible for full green resistance to AcEm2. Five-week old plants were spray-inoculated with *A. candida* and incubated at 21 °C; and then phenotyped 14 days after inoculation. Abaxial and adaxial picture of the same leaf are shown. Scale =1cm.

The *WRR4B*^Col-0/Ws-2^ paralogs also confer resistance to Ac2V in *A. thaliana* and in transgenic *B. juncea* (Cevik *et al*., 2019). However, Ws-2, which lacks *WRR4A*, exhibits full GR phenotype in adult leaves following inoculation with either Ac2V or Ac7V indicating that *WRR4B*^Ws-2^ also confers adult plant resistance to Races 2 and 7 of *A. candida* (Table **S2**).

### Selection of candidate *A. candida* CCG effectors to test for recognition by different *WRR4* paralogs

The recent PacBio based genome assembly of *A. candida* Race 2 isolate Ac2V (“Ac2VPB”) revealed that the CCG family of secreted proteins corresponds to ca. 10% of the *A. candida* secretome (Furzer *et al*., 2021). The genome of each *A. candida* race contains 60-80 CCG effector candidate genes that can vary by sequence and presence/absence polymorphism (Jouet *et al*., 2019; Furzer *et al*., 2021). This enabled identification of candidates for functional screening of CCG effectors transiently, testing for *WRR* gene-dependent HR. Such transient assays have been adopted for other oomycete pathogens like *P. infestans*, testing secreted proteins with an RxLR motif (Vleeshouwers *et al*., 2008; Vleeshouwers *et al*., 2011), and thus, identifying several recognized effector genes.

To identify CCG effector proteins that are recognized by *WRR4* paralogs, we analysed CCGs predicted from Illumina-based genome assemblies of multiple *A. candida* races (AcNc2, AcEm2, Ac7V, AcBoT and Ac2V) (McMullan *et al*., 2015; Jouet *et al*., 2019) as well as additional CCGs predicted from Ac2VPB (Furzer *et al*., 2021). We identified CCGs showing presence/absence polymorphism or pseudogenisation (Jouet *et al*., 2019; Furzer *et al*., 2021). We then selected 30 candidate CCGs to screen with *WRR4A* prioritising those either absent or pseudogenised in *WRR4A*-overcoming race AcEx1. To screen with *WRR4B*, we selected 13 candidate CCGs that are mostly conserved in Ac2V, Ac7V and AcBoT but are absent or pseudogenised in other *A. candida* races (Fig. **2a,b** and Fig. **3a**; Table **S3**).

**Figure 2.**
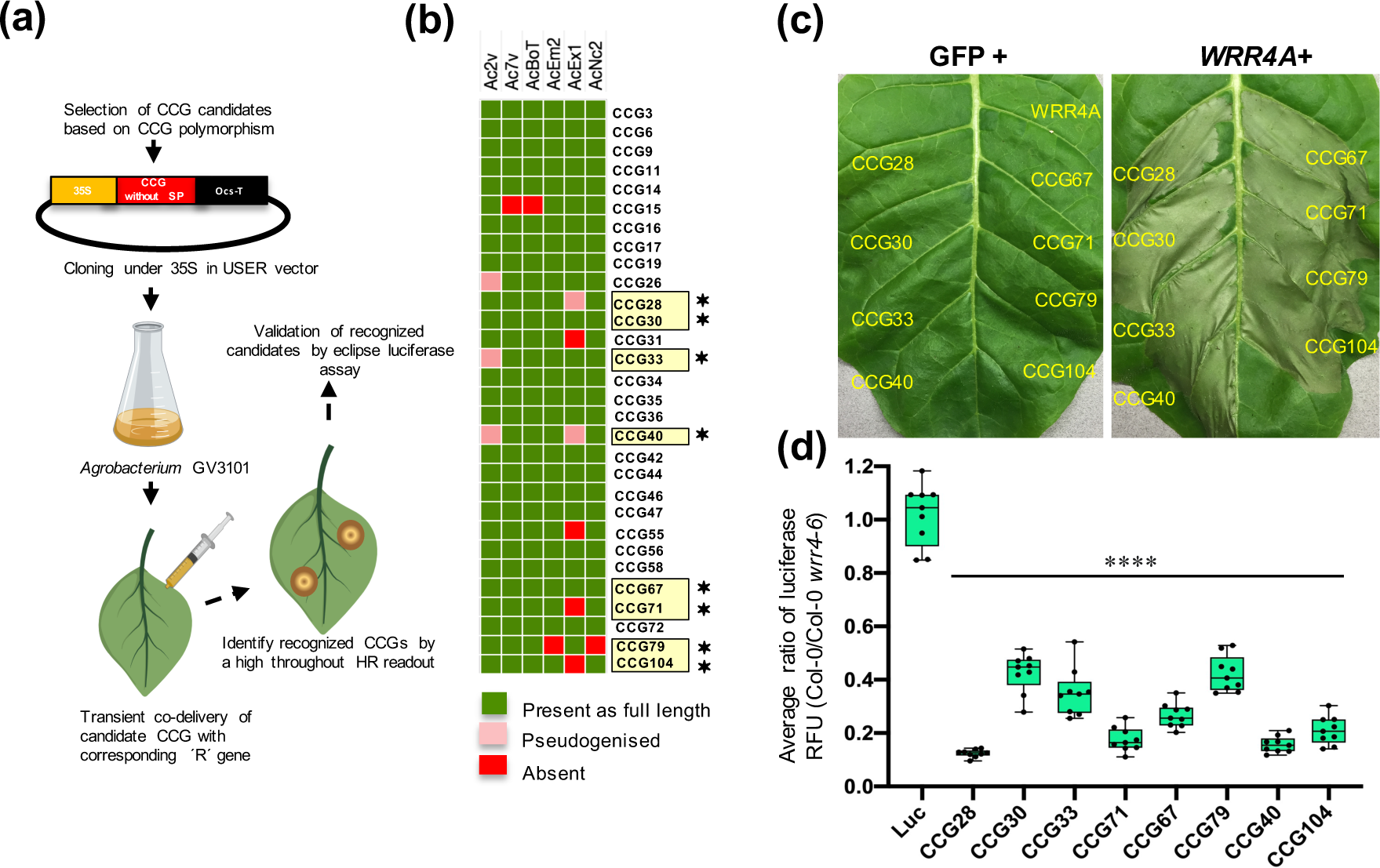
Identification of eight CCG effectors from *A. candida* that elicit hypersensitive response when co-expressed in tobacco leaves with *WRR4A* from Col-0 *A. thaliana*. **(a)** Pipeline for high-throughput screening of the CCG effectors for identification of recognized-CCGs against *WRR4A*^Col-0^ and *WRR4B*^Col-0^. The selected candidate effectors without the signal peptide (SP) were cloned into a USER expression vector under CaMV-35S promoter for Agrobacterium-mediated transient expression either in *N. tabacum* or *N. benthamiana*. Recognized CCGs were identified by an HR readout and were further validated by a luciferase eclipse assay. **(b)** The selection of candidate CCG effectors for screening against *WRR4A* was based on positive selection, presence across different *A. candida* races and/or showing the characteristic of either allelic truncation or absence in a Col-virulent isolate of Race 4 (AcEx1). The recognized CCGs against *WRR4A* identified from our screen are highlighted with a yellow box and marked with an asterisk (*). **(c)**Transient expression of candidate CCG effectors either with GFP or with *WRR4A*^Col-0^ in *N. tabacum*. The eight identified CCGs trigger HR when co-expressed with *WRR4A*^Col-0^ but not with GFP alone. *WRR4A*^Col-0^ also do not show any auto-activity when co-delivered with the GFP control. **(d)** The *WRR4A*-dependent CCG recognition correlates with the luciferase eclipse assay upon particle bombardment in *A. thaliana*. Graph showing a drop in the luciferase activity of all recognized CCGs identified by a ratio of the measured luciferase in wildtype Col-0 as compared to that Col-*wrr4a-6* mutant validating the recognition to be specific by *WRR4A*^Col-0^. Statistical significance versus Luciferase (Luc) alone (p < 0.05, one way ANOVA, Bonferroni’s multiple comparison test) is indicated by an asterisk. Error bars indicate SD (n = 9).

**Figure 3.**
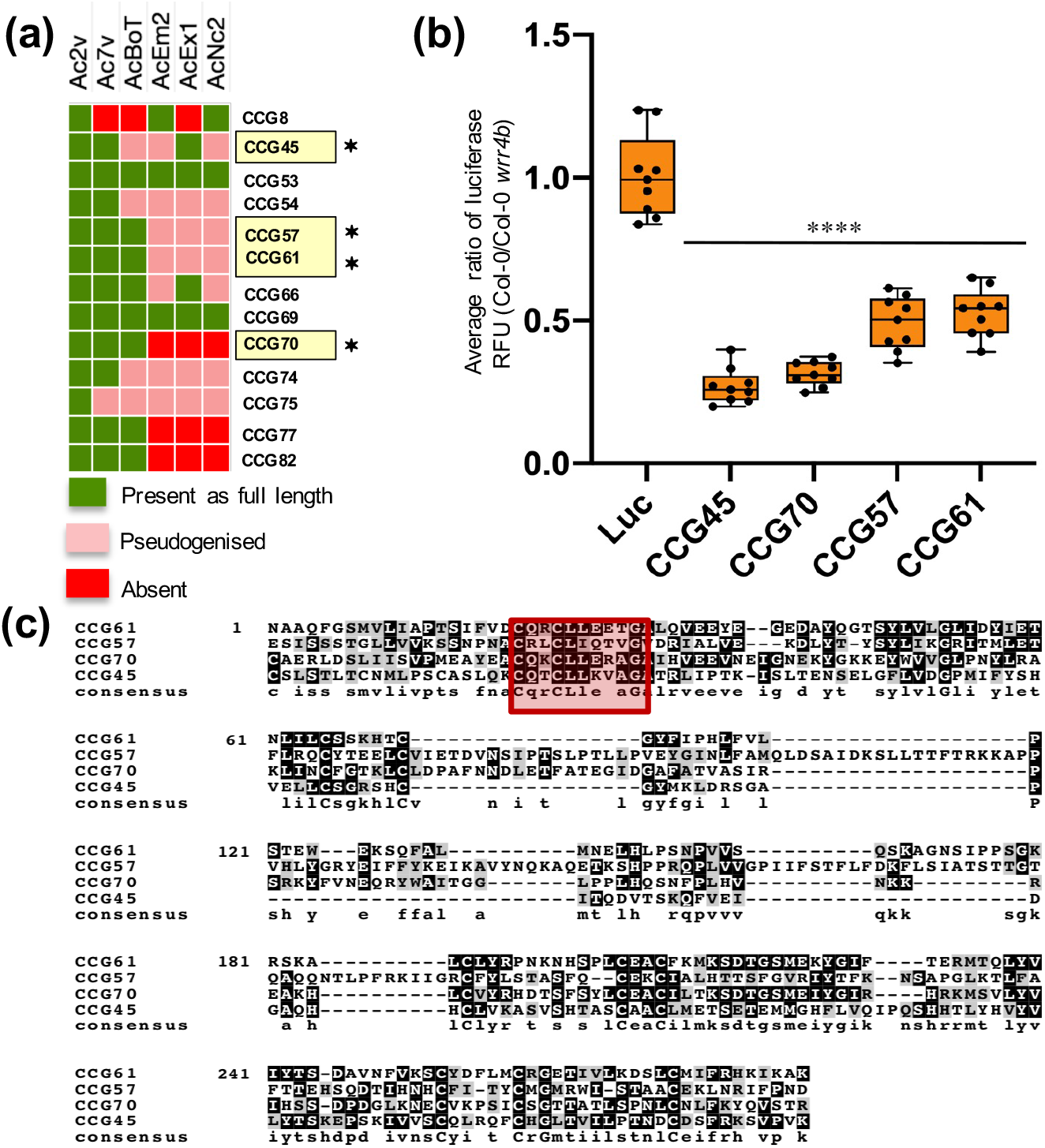
The *WRR4B*^Col-0^ paralog recognizes four additional CCG effectors in *A. candida*. **(a)** The selection of candidate CCG effectors for screening against *WRR4B*^Col-0^ based on their presence across isolates of different *A. candida* races. The recognized CCGs against *WRR4B*^Col-0^ identified from our screen are marked with an asterisk (*). **(b)** The luciferase eclipse assay upon particle bombardment in *A. thaliana*. Graph showing a drop in the luciferase activity of all recognized CCGs identified by a ratio of the measured luciferase in wildtype Col-0 as compared to a Col-*wrr4b* mutant validating the recognition to be specific by *WRR4B*^Col-0^. Statistical significance versus Luciferase (Luc) alone (p < 0.05, one way ANOVA, Bonferroni’s multiple comparison test) is indicated by an asterisk. Error bars indicate SD (n = 9). **(c)** Multiple sequence alignment of *WRR4B*-recognized CCGs which shows homology primarily in the CCG motif. CCG motif is highlighted by red box.

To test the selected CCG effectors for *WRR4A* or *WRR4B* recognition, we cloned CCG effectors into an expression vector with 35S promoter and transformed into *Agrobacterium* for infiltration of *N. tabacum* or *N. benthamiana* leaves. The transient co-delivery of CCG effector was performed either with GFP or RFP as negative control and with *WRR4A* or *WRR4B* (Cevik *et al*., 2019). Effectors that trigger an HR when co-expressed with the corresponding NLR were further validated by the luciferase eclipse assay modified from Allen *et al*., (2004) for recognition in Col-0 or in Col-0 *wrr4a* or *wrr4b* mutants (Fig. **2a**).

### *WRR4A*^*Col-0*^ confers recognition to eight different CCG effectors from *A. candida*

To screen candidate CCGs for their recognition by *WRR4A*, CCGs excluding the signal peptide were cloned from isolates of *A. candida* Race 2 (Ac2V) or Race 4 (AcEm2 or AcNc2). All of the effector alleles were co-infiltrated with 35S:*WRR4A-*expressing Agrobacterium strains (Cooper *et al*., 2008; Cevik *et al*., 2019). All of the recognized effectors and some representative non-recognized CCGs were tested by Western blot to confirm their protein expression when expressed transiently (Fig. **S1**). Among the 30 alleles we tested, eight CCGs (CCG28^Ac2V^, CCG30^AcNc2^, CCG33^AcNc2^, CCG40^AcEm2^, CCG67^AcEm2^, CCG71^AcNc2^, CCG79^Ac2V^ and CCG104^Ac2V^) elicited HR within 36-48 hours post infiltration (hpi) when co-expressed with *WRR4A* but not with GFP control. Moreover, *WRR4A* co-expressed with GFP did not show any autoactivation (Fig. **2c**).

To validate the HR observed from transient assays with the eight recognized CCGs, we carried out luciferase eclipse assays by transient delivery using particle bombardment to reveal reduced luciferase activity upon HR triggered by these recognized CCG effectors in *Arabidopsis* leaves in Col-0 compared to Col-0 *wrr4a-6* mutant. The luciferase construct was delivered alone or with recognized CCG effector and incubated in vials with water for 48 hours post-bombardment (hpb). All the CCGs recognized by *WRR4A* conferred reduced luciferase activity in comparison to the luciferase only control in Col-0 but not in Col-0 *wrr4a-6* mutant. The diminished luciferase activity in Col-0 indicates HR-dependent cell death in transformed leaf cells (Fig. **2d**). Hence, both methods revealed eight different CCGs that are recognized by *WRR4A*.

### The *WRR4A* ^*Col-0*^ paralog *WRR4B* ^*Col-0*^ recognizes four additional CCG effectors

To test recognition by *WRR4B*, we selected candidate CCGs from the CCG effectorome repertoire based on the resistance and susceptibility disease phenotypes (Table **S2**). As *WRR4B* confers resistance against *A. candida* race 2 (Ac2V), we prioritized 13 CCG candidates present mainly in crop-infecting races of *A. candida* and also CCGs present exclusively in Ac2V (Fig. **3a**), selecting the Ac2V CCG allele for these recognition assays.

Over-expression of *WRR4B* in tobacco or *N. benthamiana* shows weak autoimmunity, even when infiltrated with the GFP control (Fig. **S2 a**,**b)**. This creates a requirement for cautious interpretation of any HR phenotypes detected upon co-expression with a potentially recognized CCG effector. Hence, we used the luciferase eclipse assay for screening the set of 13 cloned CCGs for candidate *WRR4B-*recognized CCGs, and discovered four candidate *WRR4B-*recognized CCGs. We further confirmed the recognition of these four candidates (CCG45^Ac2V^, CCG57^Ac2V^, CCG61^Ac2V^ and CCG70^Ac2V^) by luciferase eclipse assay by a comparison of the luciferase activity in Col-0 and Col-0-*wrr4b* mutant. The Col-0 *wrr4b* mutant showed a higher level of luciferase when co-expressed with these 4 CCGs that is diminished in Col-0 due to HR (Fig. **3b**). This assay further confirms the *WRR4B*-specific recognition of these four CCGs from *A. candida* race Ac2V .To understand if any of these CCGs confer elevated HR after transient expression in *N. tabacum* compared to *WRR4B* alone, CCG45^Ac2V^, CCG57^Ac2V^, CCG61^Ac2V^ and CCG70^Ac2V^ were co-infiltrated with *WRR4B*. Only CCG45^Ac2V^ and CCG70^Ac2V^ showed an enhanced HR compared to the *WRR4B* infiltrated with GFP alone, indicating that these candidates are also recognized by *WRR4B* in tobacco (Fig. **S2a**). This HR varied between individual leaves. CCG57^Ac2V^ and CCG61^Ac2V^ did not show elevated HR compared to *WRR4B* co-infiltrated with GFP control (Fig. **S2a**). In order to check if the HR by CCG45^Ac2V^ and CCG70^Ac2V^ is also consistent in *N. benthamiana*, we performed co-infiltration of these CCGs with *WRR4B* in *N. benthamiana*. CCG28^Ac2V^ co-delivered with *WRR4A* was used as a positive control. Consistent with *N. tabacum* phenotype, CCG45^Ac2V^ and CCG70^Ac2V^ show an elevated HR in *N. benthamiana* when co-expressed with *WRR4B* as compared to *WRR4B* alone (Fig. **S2b, c**). The CCG motif region is the most conserved part of different *WRR4B*-recognized CCGs (Fig. **3c**). We conclude that *WRR4B* specifically recognizes four CCGs from *A. candida* that are distinct from the CCGs that are recognized by *WRR4A*.

To further elucidate the region of CCG that is recognized, we investigated the *WRR4A*-CCG interaction, as *WRR4A* does not show any autoimmune phenotype and also recognizes diverse CCGs across different *A. candida* clades (Furzer *et al*., 2021).

### Recognition of CCGs by *WRR4A*^Col-0^ requires the N-terminal portion of the protein but does not occur via the CCG motif

To investigate the mechanism of *WRR4A* activation by CCGs, we tested a series of CCG effector deletions. We selected CCG28 for this analysis because it triggers the strongest *WRR4A*-dependent HR, with a response visible at 36 hpi. Similar to *WRR4B-*recognized CCGs, as the CCG motif region was the only part of the CCG protein that showed homology across different *WRR4A* recognized CCGs, we tested the role of the N-terminal region of CCG for recognition by *WRR4A*. A full-length version of CCG28 without its secretion signal triggers strong *WRR4A*-dependent HR. However, deletion of amino acids 28-40 in the N-terminal region after the secretion signal abolishes this recognition (Fig. **4a, b**).

**Figure 4.**
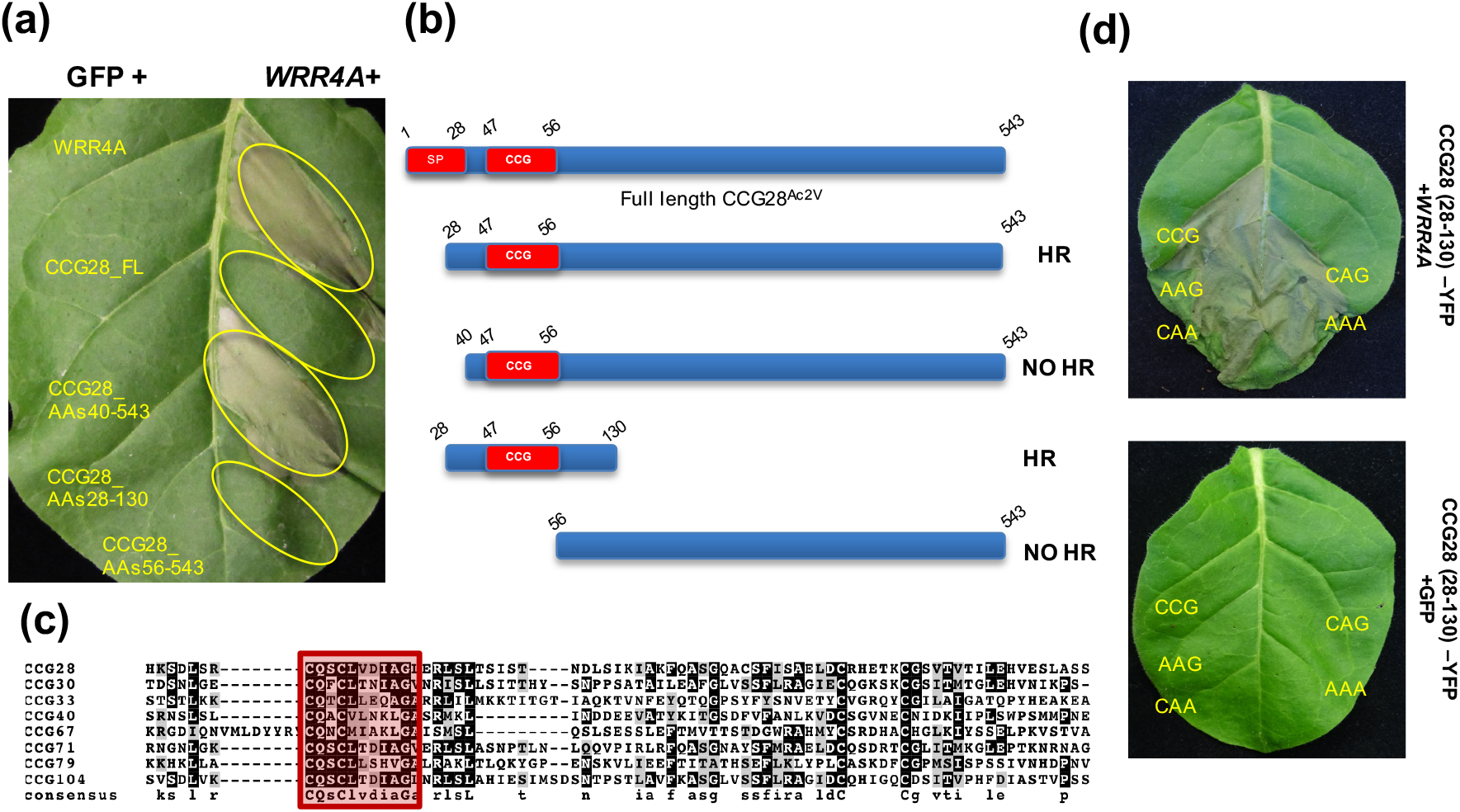
The N-Terminal Portion of CCG effectors is sufficient for recognition by *WRR4A*^Col-0^ and occurs independent of the CCG motif. **(a)**Transient expression of different truncated versions of CCG28 either with GFP or with *WRR4A*^Col-0^ in *N. tabacum*. The N-terminal region that corresponds to aa 28-130 is mainly indispensable for *WRR4A*^Col-0^ recognition. **(b)** Schematic representation of all the truncations infiltrated in Fig. 4a, showing either a HR or a No-HR phenotype. **(c)** The alignment of protein sequences of N -terminal region of CCGs after signal peptide cleavage, up to amino acid 130 in all *WRR4A*^Col-0^ recognized CCGs. CCG motif is highlighted by red box. **(d)** Mutations in CCG motif with CCG changed to AAG, CAA, CAG and AAA do not impair *WRR4A*^Col-0^ recognition and HR. *N. tabacum* leaf was infiltrated to express wild-type or mutated variants of CCG motif in CCG28^aa28-130^ -YFP. All versions with wild-type as well as mutated CCG motif trigger HR, which was assessed at 48 hpi. Photographs are representative of three consistent replicates.

To define the minimal N-terminal region of CCG28 that is recognized, we made C-terminal deletions of CCG28^Ac2V^ and tested their recognition using transient assays in *N. tabacum* leaves (Fig. **S3**). A truncation of CCG28^Ac2V^ that includes the first 100 amino acids after the signal peptide site (CCG28^28-130^), including the CCG motif, is more strongly recognized by *WRR4A* than full length CCG28, triggering an elevated HR at 36 hpi when transiently co-expressed in *N. tabacum* (Fig. **4a, b**; Fig. **S3**). This also includes a common feature with a pair of cysteines located ∼50 amino acids after the CCG motif. In contrast, a C-terminal region of CCG28 without the CCG motif, which corresponds to aa 56-543 abolishes recognition when co-expressed with *WRR4A* (Fig. **4a, b**; Fig. **S4**). Hence, we conclude, that the truncated N-terminal part of CCG28 is indispensable for *WRR4A* recognition. Further deletion of amino acids 28-35 at the N-terminus also compromises recognition (Fig. **S3** and **S5**). To define the shortest region of CCG28 that is recognized, we further narrowed the recognition region to 50 amino acids, corresponding to CCG28^28-78^. However, only YFP-tagged versions of this shortest region activate HR (Fig. **S3** and **S5**), likely indicating that untagged versions are insufficiently stable for their interaction with *WRR4A* to trigger a cell death phenotype when expressed transiently.

We next tested whether *WRR4A* recognition of all the CCG candidates involves their N-termini. A truncation analysis was carried out for all the eight recognized CCGs by *WRR4A*. Upon transient expression, C-terminally truncated versions of the other seven recognized CCGs also trigger an HR phenotype with *WRR4A* in *N. tabacum* (Fig. **S6a**). These data suggest that the N-terminal portion of all the *Avr-WRR4A* is sufficient for recognizability. *WRR4A* recognition of CCGs requires an intact Walker A (P-loop) (Schreiber *et al*., 2016) because a mutation in this P-loop of *WRR4A* with a change from K220 >L220 abolishes the HR when *WRR4A* is co-infiltrated with the recognized CCGs in transient assays (Fig. **S6b**).

We next examined whether the previously defined CCG motif, that shows maximum homology across all recognized CCGs against *WRR4A* mediates effector recognizability (Fig. **4c**). To this end, the truncated CCG28^aa28-130^, which is strongly recognized by *WRR4A* was used. Mutant versions of CCG28^aa28-130^-YFP carrying a mutation in the CCG motif (where CCG exchanged to AAG, CAA, CAG and AAA) were generated and tested for recognition in transient assays by co-infiltration with *WRR4A* in *N. tabacum*. These mutated versions were still strongly recognized by *WRR4A* with an HR phenotype indistinguishable from the truncated version with authentic CCG motif (Fig. **4d**). These data suggest that *WRR4A* -CCG recognition does not occur via the CCG motif. Conceivably a structural similarity in the N-terminal portion of these recognized CCGs might be responsible for their detection by *WRR4A*. Consistent with this, CCG30, but not its close paralog CCG16, is recognized by *WRR4A* (Fig. **S7**).

### Allelic variation and expression polymorphism of recognized *WRR4A* CCGs

Next, we confirmed the allelic status of the identified recognized CCG candidates across all the sequenced races of *A. candida* from the available Illumina assemblies of different *A. candida* races (McMullan *et al*., 2015) and the recent PacBio assembly of *B. juncea*-infecting race Ac2V (Furzer *et al*., 2021). Consistent with the resistance phenotypes in Col-0 and Col-*wrr4a* mutants, the recognized CCGs are present across different *A. candida* races. CCG33 and CCG40 are pseudogenised in the *B. juncea*-infecting race Ac2V (Fig. **2b**). Several of the recognized alleles from these CCG candidates, show polymorphism in AcEx1 (a *WRR4* resistance-breaking pathotype of Race 4). Due to an early stop codon, CCG28^AcEx1^ shows a protein length of only 226 amino acids as compared to the full-length alleles from other races (Fig. **5a**). Another isolate of this pathotype, AcCarlisle collected on *A. thaliana*, shows the same early stop codon in CCG28 and virulence on *Arabidopsis* Col-0. The AcEx1 CCG28 allele is weakly recognized compared to full-length alleles from *A. candida* Races 2, 4 and 7 (Ac2V, AcEm2 and Ac7V) all of which trigger an early *WRR4A*-dependent HR at 36 hpi (Fig. **5b**). Additionally, the *WRR4A-*recognized CCG71 and CCG104 are absent from AcEx1 (Fig. **2b**) suggesting that absence of these effectors might contribute to evasion of *WRR4A* mediated resistance. On the other hand, the CCG effectors recognized by *WRR4B* showed a strong presence/absence polymorphism and are primarily present in the crop-infecting races which are known to be resisted by *WRR4B* (Fig. **3a**).

**Figure 5.**
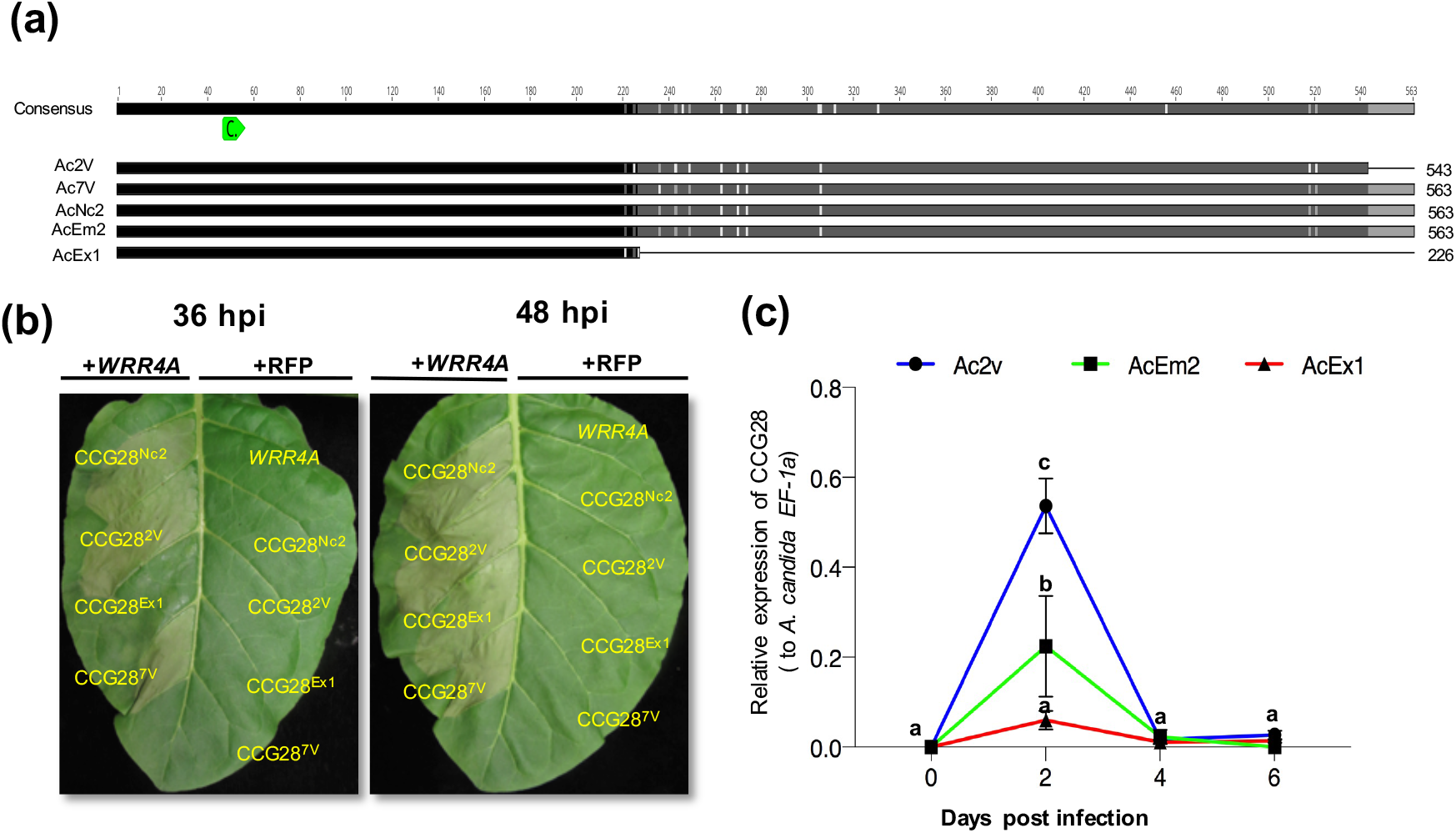
Differential recognition of CCG28 alleles across different *A. candida* races. **(a)** Schematic representation of the amino acid sequence alignment of CCG28 alleles from *A. candida* Race 2 (Ac2V), Race7 (Ac7v) and Race 4 (AcNc2, AcEm2 and AcEx1). The black region indicates conserved amino acid sequence across all races, whereas the grey region indicates variation. White bars in the alleles indicate the polymorphic sites where there is no predominant residue, and grey bars indicate polymorphic sites where there is a predominant residue. The numbers next to the alleles refer to the total protein length. Figure drawn to scale. The CCG motif is indicated in green with a C. **(b)** Quantitative recognition of different CCG28 alleles in *N. tabacum* transient assay. The different CCG28 alleles were co-infiltrated with *WRR4A*^Col-0^ into *N. tabacum* leaves. CCG28 shows recognition with an enhanced HR at 36 hpi. However, the AcEx1 CCG28 allele with a premature stop codon shows a delayed HR at 48 hpi, consistent with weak recognition of the CCG28 allele in an *A. candida* Race 4 isolate that overcomes the *WRR4A*^Col-0^ recognition. **(c)** Expression analysis of CCG28 (relative to *A. candida EF-1a*) in Race 2 (Ac2v) and Race 4 (AcEm2 and AcEx1), showed that the AcEx1 truncated allele of CCG28 is weakly expressed, likely contributing to virulence on *WRR4A*^Col-0^. Different letters indicate statistically significant differences between the different alleles tested (2 way ANOVA, Bonferroni’s multiple comparison test, p < 0.05). Error bars represent SD.

To determine the extent to which these *WRR4A-*and *WRR4B*-recognized CCGs are expressed *in planta*, we assessed their expression profiles from the RNA-Seq data obtained over the consecutive time-points during serial infection stages of *A. candida* Race Ac2V (Furzer *et al*., 2021). *WRR4A-*and *WRR4B*-recognized CCGs in Ac2V show *in planta* induction during different colonization stages. These CCGs are induced specifically at different colonization time-points of 2, 4 and 6 dpi (Fig. **S8a**) consistent with a role in virulence.

To investigate the expression patterns of different alleles of the *WRR4A*-recognized CCGs, we performed a qRT-PCR expression profiling of the *A. candida* races Ac2V, AcEm2 and AcEx1 growing on Ws*-eds1* plants (Fig. **S8b**). These data showed different CCG effectors increase in expression during infection. Intriguingly, CCG28 is primarily induced at 2 dpi. The Ac2V allele of CCG28 is most highly expressed followed by the AcEm2 allele. Notably, the pseudogenised allele from AcEx1 shows least expression as compared to the other alleles (Fig. **5c**). The expression profiles of other recognized *WRR4A*-recognized CCGs follow a similar trend. The Ac2V allele is among the most highly expressed and AcEx1 shows the lowest (Fig. **S8c**). From these results, we conclude that the partial susceptibility of Col-0 plants we observe with AcEx1 is both due to reduced expression or due to absence of the *WRR4A*-recognized CCGs. In summary, these data suggest for *WRR4*-recognized CCGs, there exists expression polymorphism as well as presence-absence polymorphism that can enable evasion of resistance, especially when combined with the well-documented suppression of resistance upon *Albugo* infection.

### AcEx1 has the capacity to suppress *WRR4A*-mediated resistance

Previous studies have shown that *A. candida* infection leads to strong immune suppression enabling co-infection by otherwise avirulent races, permitting sexual exchange and recombination (Cooper *et al*., 2008; McMullan *et al*., 2015). To test if AcEx1 can suppress *WRR4A* mediated resistance, we performed sequential inoculation experiments, on *Arabidopsis* accessions with and without *WRR4A*, monitoring *A. candida* races using race-specific genomic regions as a readout for race-specific PCR markers. Race-specific PCR verifies that AcNc2 can grow on Ws-2 which does not have *WRR4A* but not on RIL CW20 (which carries Col-5 alleles of *WRR4A* and *WRR4B* as the only *WRR* genes). However, pre-inoculated plants of CW20 colonized by the *WRR4A* resistance breaking race AcEx1 lose resistance in CW20 leaves towards AcNc2 (Fig. **S9**). Therefore, AcEx1 not only suppresses *WRR4A* mediated recognition but also enables other races to grow, that would otherwise be resisted. These data suggest that AcEx1 is able to overcome *WRR4A* resistance by virtue of weak recognizability due to polymorphisms in its repertoire of *WRR4A* recognized CCG alleles, together with strong ability for immunosuppression of *WRR4A*-mediated resistance.

## DISCUSSION

Oomycetes in the Peronosporales (*Phytophthora, Pythium* and downy mildew species) are destructive pathogens. Host colonization requires effectors which carry a signal peptide and a positionally constrained RxLR motif (Win *et al*., 2006). A specific effector is often detected in a resistant host by a matching NLR immune receptor, often encoded by a resistance (*R*-) gene, that activates a defense response that thwarts pathogen success (Armstrong *et al*., 2005; Rehmany *et al*., 2005). The Irish potato famine pathogen *P. infestans*, RxLR effectors *Avrblb2* and *Avrvnt1* are recognized by *Rpi-blb2* and *Rpi-vnt1* (Oh *et al*., 2009; Pais *et al*., 2018). Identifying such avirulence (*Avr*) determinants in the pathogen is important for improving strategies to develop disease resistant crops.

The genomes of oomycetes in the *Albuginales*, such as *A. laibachii* and *A. candida*, are not enriched for RxLR proteins but instead contain a family of secreted proteins with a ’CHxC’ (now CCG) motif that contributes to translocation (Kemen *et al*., 2011; Furzer *et al*., 2021). These proteins show signatures of diversifying selection, with high ratios of non-synonymous to synonymous mutations, similar to RxLR effectors (Rehmany *et al*., 2005; Asai *et al*., 2018). The Ac2VPB assembly of the Brassica pathogen *A. candida* Race 2 has enabled refinement of CCG effector annotation (Furzer *et al*., 2021). CCG gene repertoires are present in all sequenced *A. candida* races (McMullan *et al*., 2015). A combined phylogeny comparison between *A. laibachii* and *A. candida* shows an expansion of CCGs from ∼30 in *A. laibachii* to around ∼100 in *A. candida*, with several *A. candida*-specific clades suggesting expansion and reshuffling of CCG effector repertoires to adapt to different hosts (Furzer *et al*., 2021). Moreover, the CCG repertoire displays elevated rates of pseudogenisation and presence/absence polymorphisms, consistent with selection for diversity while maintaining virulence functions.

In this study, we used the CCG effectoromes from different *A. candida* races, enabling us to screen selected CCG effectors for *WRR4* recognition. This revealed eight *WRR4A-*recognized CCGs and four *WRR4B*-recognized CCGs. The allelic comparison approach helped us to select CCG candidates based on their presence/absence polymorphism in *A. candida* races that overcome resistance. The eight identified *WRR4A-*recognized CCGs when co-expressed with *WRR4A* show an HR at 48 hpi resulting in complete cell-death. A genetic validation of this recognition by luciferase eclipse assay after particle bombardment shows that the identified CCG effectors are specifically recognized by *WRR4A*. A similar approach based on allelic comparison allowed us to screen and identify four additional *WRR4B*-recognized CCGs. As *WRR4B* shows an autoimmune phenotype in transient expression experiments, the *N. tabacum* HR assay was more difficult to interpret than with *WRR4A*. The luciferase eclipse assay enabled us to identify four *WRR4B*-recognized CCGs; CCG45, CCG70, CCG57 and CCG61. However, only CCG45 and CCG70, which show the strongest phenotype in the luciferase eclipse assay show an enhanced HR when co-expressed transiently with *WRR4B* in *N. tabacum* or *N. benthamiana*. We conclude that *WRR4A* and *WRR4B* each recognize multiple and distinct CCG effectors from *A. candida*. Multiple recognition of effectors is known for some other NLRs that function in pairs, such as *Arabidopsis* RPS4/RRS1 (Sarris *et al*., 2015; Guo *et al*., 2020) and rice RGA4/RGA5 (Cesari *et al*., 2013). The *WRR4B* autoimmune phenotype hindered further analysis, and we further investigated the *WRR4A*-CCG interaction to define recognition requirements in more detail.

Analysis of the recognition mechanism of multiple CCGs by *WRR4A* revealed that the N-terminal portion of all *WRR4A-*recognized CCGs is sufficient for recognizability. Truncation analysis of all these CCGs shows the N terminal 12 amino acids after the secretion signal are required to trigger HR. Mutations in the CCG motif suggest that the CCG motif itself is not recognized, as mutant alleles are still recognizable. Hence, the exact role of the CCG motif and its significance in *A. candida* infection is still unclear. The recognized CCGs fall into different clades in the CCG phylogeny (Furzer *et al*., 2021). Hence, we speculate that there likely exists a structural similarity among these recognized effectors that enables their recognition. This is also suggested by the observation that CCG16, a close paralog of the *WRR4A*-recognized CCG30 is not recognized despite showing a higher homology to CCG30 than some other *WRR4A-*recognized CCGs.

This study provides evidence that *WRR4A* and *WRR4B* at the WRR4 locus in *A. thaliana* Col-0 have the capacity to recognize multiple effectors from *A. candida* likely making them highly effective for immune activation upon pathogen attack. This potentially explains the effectiveness of this resistance against many races. Interestingly, both *WRR4A* and *WRR4B* have a post-LRR C-JID domain which was recently shown to physically interact with effectors by TNLs RPP1 and Roq1 during resistosome activation (Ma *et al*., 2020; Martin *et al*., 2020). Hence, it will be intriguing to understand whether *WRR4*-CCG interaction is mediated by a similar mechanism. For *P. infestans* RxLR effectors, the C-terminal post-RxLR domain carries their effector activities (Kamoun, 2006; Kamoun, 2007). It is unknown how CCG proteins contribute to pathogen virulence and is a topic of further investigation. AcEx1 is capable of overcoming *WRR4A-*and *WRR4B*-mediated resistance, though on Col-0, but not Col-0 *wrr4A*, it activates chlorosis and thus is still weakly recognized. Its growth on Col-0 is due to the loss of some of the recognized *A. candida* effectors, and reduced expression of others.

Additionally, AcEx1 suppresses *WRR4*-mediated immunity against AcNc2, consistent with the hypothesis that *Arabidopsis* susceptibility to a specific *A. candida* race is determined by a balance of effector detection and effector-mediated suppression of defense. The partial susceptibility of Col-0 plants with AcEx1 is potentially explained by the reduced expression (measured by expression profiling) of otherwise detectable CCG effectors in this isolate. We suggest that strong immunosuppression of this weak recognition enables AcEx1 to colonize Col-0. Therefore, we propose that further study of *WRR4A-*and *WRR4B-*recognized CCGs might reveal insights into the colonization biology of *A. candida*.

Resistance to AcEx1 that maps to the *WRR4* locus has been identified in other accessions of *Arabidopsis* (Fairhead, 2016; Castel *et al*., 2021). The *WRR4A* alleles in HR-5 and Oy-0 have an extended C-terminus, that confers AcEx1 resistance by recognizing alternative CCG effectors. The identification of *WRR4A-*and *WRR4B-*recognized CCGs will further enable understanding of the recognition mechanism at the structural level. Thus, this work contributes new insights into effector biology in obligate biotrophs, and will help inform provision of durable resistance in Brassicaceae crops by pyramiding or transgene stacking of different *WRR4* paralogs which recognize diverse repertoires of CCG effectors.

## Supporting information

Suppmental Figures

## Author Contributions

A.R., V.C. and J.D.G.J conceptualized and designed the research. A.R., V.C. and K.B. S.F. conducted all experiments. A.R., V.C., K.B., O.J.F. and S.F. performed the data analysis. M.H.B. and E.H gave critical intellectual input and provided material for this work. A.R., V.C. and J.D.G.J wrote the manuscript with input from all co-authors. All authors helped editing and finalizing the manuscript.

## Acknowledgements

A.R. acknowledges support by EMBO LTF (ALTF-842-2015). V.C., O.J.F. and S.F. were supported by Biotechnology and Biological Sciences Research Council (BBSRC) grant BB/L011646/1. K. B. and J.D.G.J. were supported in part by ERC Advanced Investigator grant to JDGJ ‘ALBUGON’ Project ID 233376. Research in the Jones Lab is supported by the Gatsby Foundation (UK) and BBSRC. We thank Shihomi Uzuhashi for her help in initial screening of the *Albugo* effectors in tobacco.

