## Supplementary material for "The Arabidopsis *WRR4A* and *WRR4B* paralogous NLR proteins both confer recognition of multiple *Albugo candida* effectors": Suppmental Figures

The following Supporting Information is available for this article:

**Fig. S1 Confirmation of expression of *WRR4A* recognized and representative non-recognized CCGs.**

Western Blot analysis of Flag-His tagged recognized and non-recognized CCGs. The CCG effectors were expressed under 35S promoter in *N. benthamiana* and leaf samples were taken 3 dpi. Asterisk indicates expected protein size. Loading control is shown by a Ponceau stained gel.

**Fig. S2 *WRR4B* shows an enhanced HR with $CCG45^{Ac2V}$ and $CCG70^{Ac2V}$**

**(a)**The *WRR4A* paralog *WRR4B* of Col-0 *A. thaliana* exhibit enhanced HR following co-expression with $CCG45^{Ac2V}$ and $CCG70^{Ac2V}$ in *N. tabacum.* Transient expression of candidate CCG effectors either with GFP or with *WRR4B* in *N. tabacum*. Two CCGs, $CCG45^{Ac2V}$ and $CCG70^{Ac2V}$ triggers an enhanced HR phenotype when co-expressed with *WRR4B* as compared to a weaker autoimmune phenotype when expressed with GFP. $CCG57^{Ac2V}$ and $CCG61^{Ac2V}$ do not show a stronger HR response as compared to the negative control with an autoimmune readout.

**(b)** Cell death observed in *N. benthamiana* leaves after expression of the CCG candidate $CCG45^{Ac2V}$ and $CCG70^{Ac2V}$. $CCG28^{Ac2V}$ co-infiltrated with *WRR4A* was used as a positive control. *N. benthamiana* leaf panels were photographed at four days after *Agrobacterium* infiltration.

**(c)** Violin plots showing cell death intensity scored as an HR index based on three independent experiments

**Fig. S3 CCG28 recognition is mediated by the N-terminal portion**

Schematic diagram illustrating the different truncations generated for the CCG28 effector to define the minimal N-terminal region that is recognized by *WRR4A* upon transient infiltration in *N. tabacum*. The HR or No HR phenotype observed for the individual tested truncated version is indicated.

**Fig. S4 CCG28 N-terminal portion of 100AA is recognized by *WRR4A***

Transient expression of different truncations of CCG28 effector either with mRFP or with *WRR4A* in *N. tabacum*. A truncation of CCG28 that includes the first 100 amino acids after the signal peptide site ($CCG28^{28\text{-}130}$), including the CCG motif, is sufficient for recognition by *WRR4A* when transiently co-expressed in *N. tabacum.* Moreover, a further deletion narrowing the recognition region to 50 amino acids, corresponding to $CCG28^{28\text{-}78}$ is also recognized. However, only YFP-tagged versions of this shortest region activate HR. In contrast, a C-terminal region of CCG28 without the CCG motif, which corresponds to aa 56-543 abolishes recognition when co-expressed with *WRR4A.*

**Fig. S5 Truncations in CCG28 showing the N-terminal portion being recognized by *WRR4A***

A rigorous recognition analysis on the truncated $CCG28^{28\text{-}130}$ shows an indispensable role of amino acids (AAs) 28-33 to be essential for an early recognition at 36hpi. Truncated versions of CCG28 that do not have AA 28-33 are delayed in recognition and recognized after around 60hpi.

**Fig. S6 CCG N-terminal portion is sufficient for recognition by *WRR4A* and is dependent on intact P-loop motif**

**(a)** Transient expression of different truncated versions of all *WRR4A* recognized CCGs either with GFP or with *WRR4A* in *N. tabacum*. The N-terminal region of all recognized CCGs if sufficient for *WRR4A* recognition.

**(b)** *WRR4A* mediated CCG recognition is dependent on intact Walker A (P-loop) motif.

**Fig. S7 *WRR4A* recognizes N-Terminal portion of CCG30 but not close paralog CCG16**

The role CCG motif is dispensable for recognition by *WRR4A.* The alignment of protein sequences of the two paralogs CCG30 and CCG16 which shows a high identity in their sequence but CCG16 is not recognized by *WRR4A.* The red triangles indicate the minimal recognized region for CCG30.

**Fig. S8 Expression profiling of recognized CCGs by RNASeq and RT-qPCR analysis**

(a) Expression profiles of all the WRR*4A*- and *WRR4B*-recognized CCGs from the RNA-Seq data obtained over the consecutive time-points during serial infection stages of *A. candida* Race Ac2V. The recognized CCGs show a clear *in planta* expression suggesting their role during plant colonization.

(b) Disease phenotypes on ws-*eds1 A. thaliana* plants upon infection *A. candida* races Ac2V, AcEm2 and AcEx1.

(c) Transcript levels of the different alleles of *WRR4A* recognized CCGs in *ws-eds1* infected plants at different timepoints after inoculation. Expression was normalized to the Ac*EF1a* gene. Different letters indicate statistically significant differences between the different alleles tested (2 way ANOVA, Bonferroni´s multiple comparison test, $p < 0.05$). Error bars represent SD.

**Fig. S9 AcEx1 can suppress *WRR4A* mediated resistance**

Co-inoculation of AcEx1 and AcNc2 onto CW20 (*WRR4*$^{\text{Col-0}}$) and Ws-2 (*wrr4*) reveals AcEx1 to suppress WRR4 mediated immunity to AcNc2 (observed as AcNc2-specific amplicon present in CW20 pre-inoculated with AcEx1).

**Table S1** Primers and plasmids used in this study

**Table S2** Resistance and susceptibility of adult leaves of *Arabidopsis* lines to various *Albugo candida* races

**Table S3** CCG protein NCBI Accession numbers

### Fig. S1

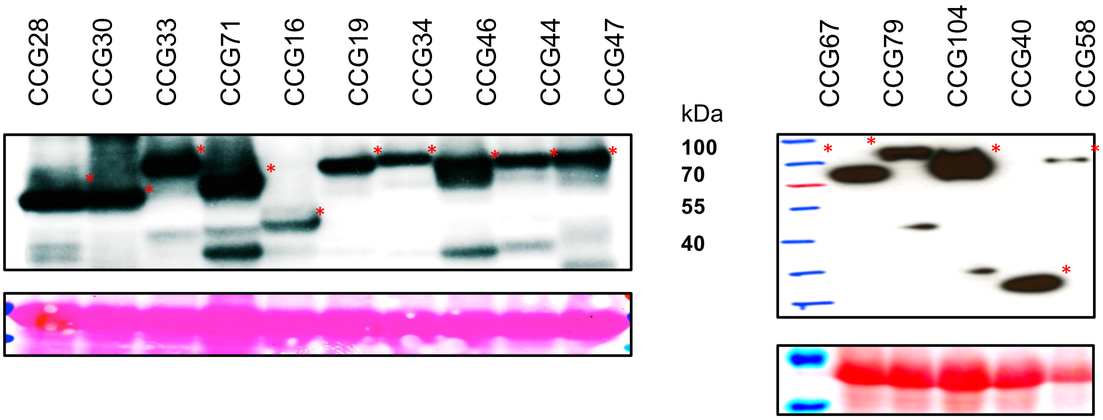


**Fig. S1 Confirmation of expression of *WRR4A* recognized and representative non-recognized CCGs.** Western Blot analysis of Flag-His tagged recognized and non-recognized CCGs. The CCG effectors were expressed under 35S promoter in *N. benthamiana* and leaf samples were taken 3 dpi. Asterisk indicates expected protein size. Loading control is shown by a Ponceau stained gel.

#### Fig. S2

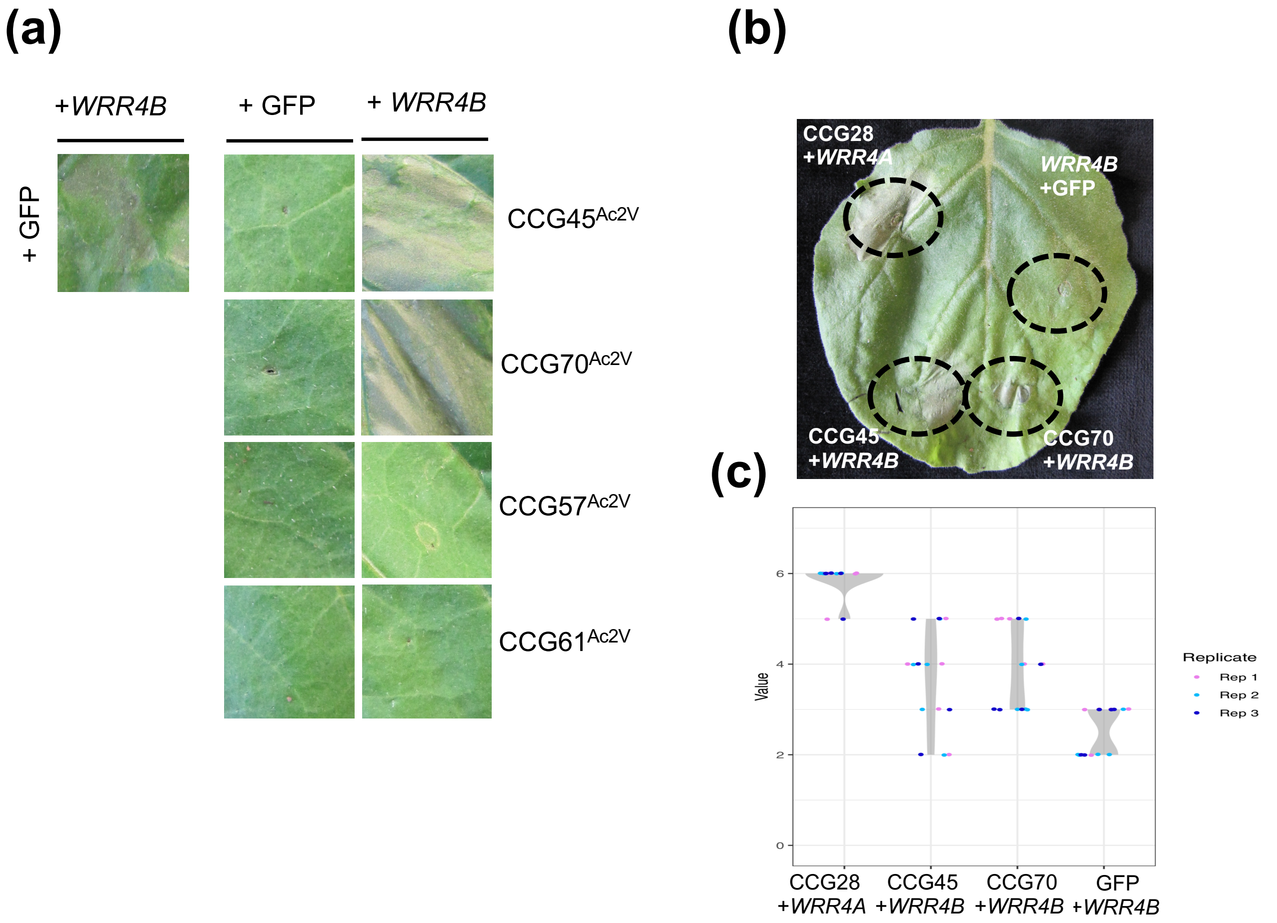


**Fig. S2 *WRR4B* shows an enhanced HR with CCG45$^{Ac2V}$ and CCG70$^{Ac2V}$**

**(a)**The *WRR4A* paralog *WRR4B* of Col-0 *A. thaliana* exhibit enhanced HR following co-expression with CCG45$^{Ac2V}$ and CCG70$^{Ac2V}$ in *N. tabacum*. Transient expression of candidate CCG effectors either with GFP or with *WRR4B* in *N. tabacum*. Two CCGs, CCG45$^{Ac2V}$ and CCG70$^{Ac2V}$ triggers an enhanced HR phenotype when co-expressed with *WRR4B* as compared to a weaker autoimmune phenotype when expressed with GFP. CCG57$^{Ac2V}$ and CCG61$^{Ac2V}$ do not show a stronger HR response as compared to the negative control with an autoimmune readout.

**(b)** Cell death observed in *N. benthamiana* leaves after expression of the CCG candidate CCG45$^{Ac2V}$ and CCG70$^{Ac2V}$. CCG28$^{Ac2V}$ co-infiltrated with *WRR4A* was used as a positive control. *N. benthamiana* leaf panels were photographed at four days after *Agrobacterium* infiltration.

**(c)** Violin plots showing cell death intensity scored as an HR index based on three independent experiments

### Fig. S3

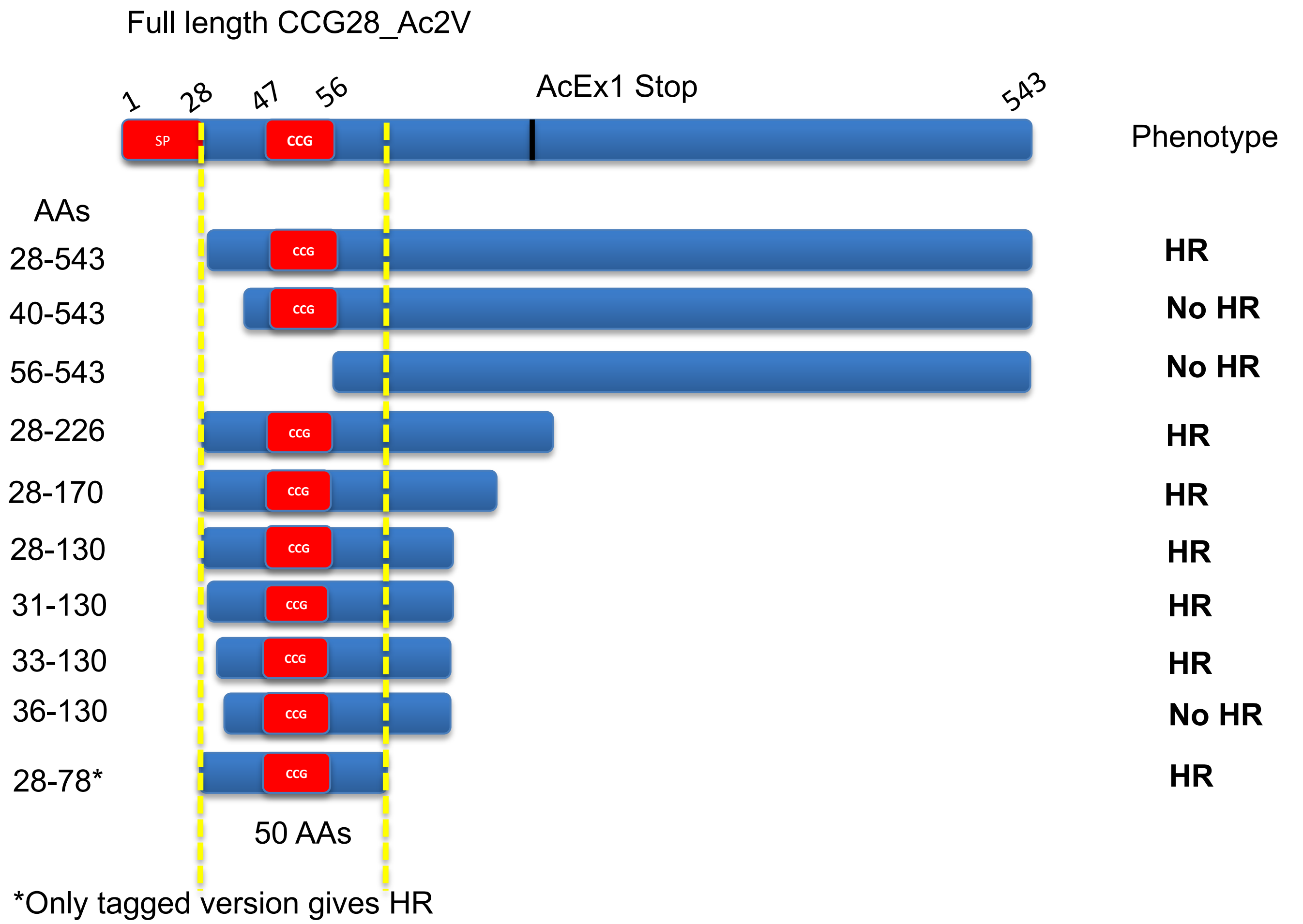


**Fig. S3 CCG28 recognition is mediated by the N-terminal portion**

Schematic diagram illustrating the different truncations generated for the CCG28 effector to define the minimal N-terminal region that is recognized by *WRR4A* upon transient infiltration in *N. tabacum*. The HR or No HR phenotype observed for the individual tested truncated version is indicated.

### Fig. S4

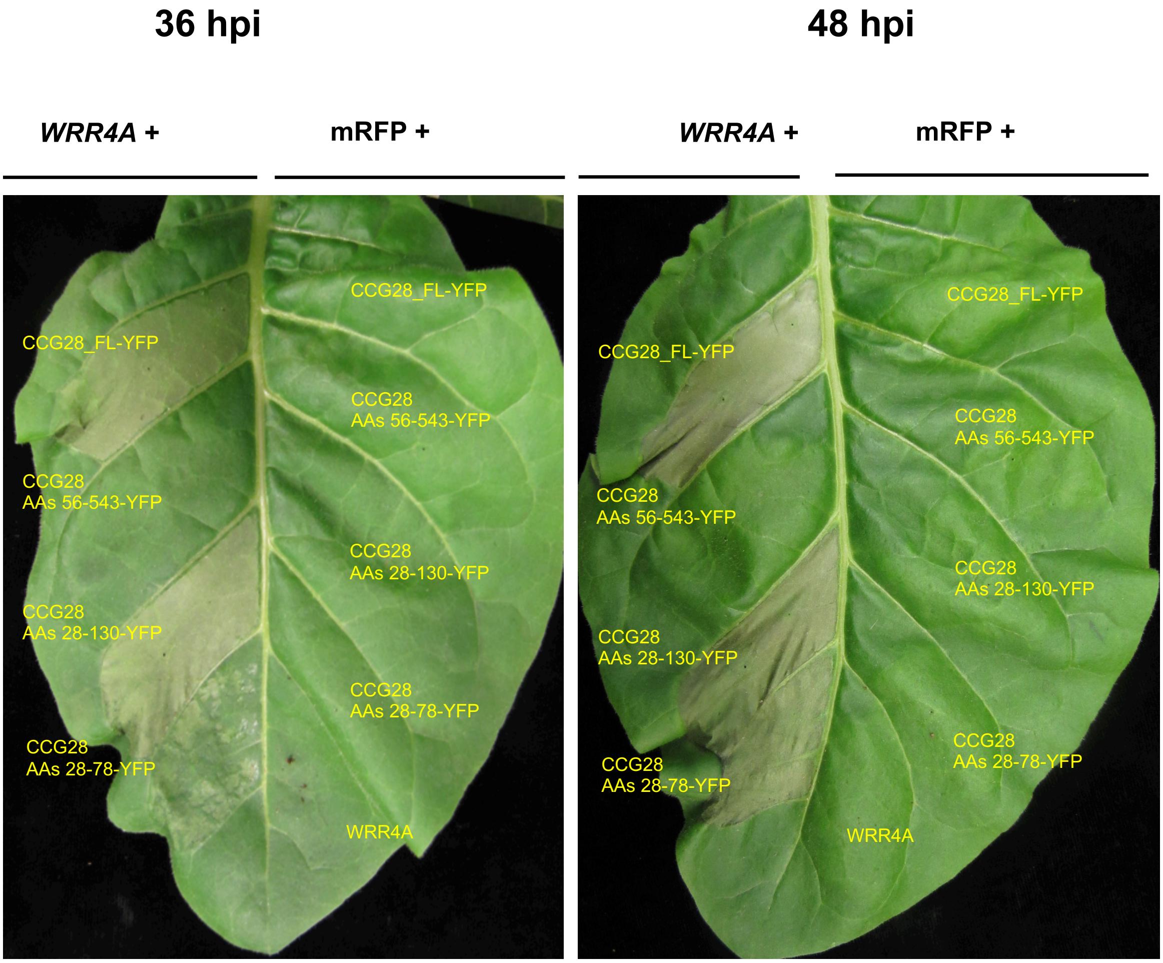


**Fig. S4 CCG28 N-terminal portion of 100AA is recognized by *WRR4A***

Transient expression of different truncations of CCG28 effector either with mRFP or with *WRR4A* in *N. tabacum*. A truncation of CCG28 that includes the first 100 amino acids after the signal peptide site ($CCG28^{28\text{-}130}$), including the CCG motif, is sufficient for recognition by *WRR4A* when transiently co-expressed in *N. tabacum*. Moreover, a further deletion narrowing the recognition region to 50 amino acids, corresponding to $CCG28^{28\text{-}78}$ is also recognized. However, only YFP-tagged versions of this shortest region activate HR. In contrast, a C-terminal region of CCG28 without the CCG motif, which corresponds to aa 56-543 abolishes recognition when co-expressed with *WRR4A*.

### Fig. S5

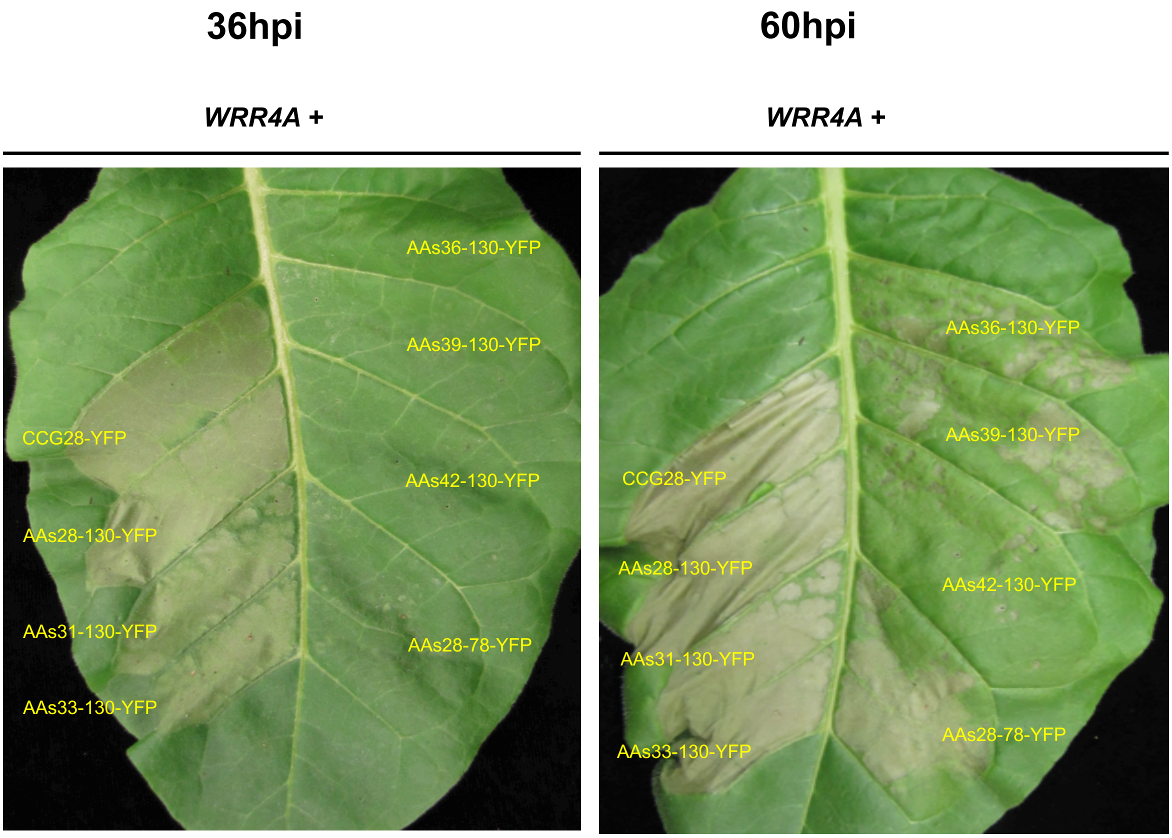


**Fig. S5 Truncations in CCG28 showing the N-terminal portion being recognized by *WRR4A***
A rigorous recognition analysis on the truncated CCG28$^{28\text{-}130}$ shows an indispensable role of amino acids (AAs) 28-33 to be essential for an early recognition at 36hpi. Truncated versions of CCG28 that do not have AA 28-33 are delayed in recognition and recognized after around 60hpi.

#### Fig. S6

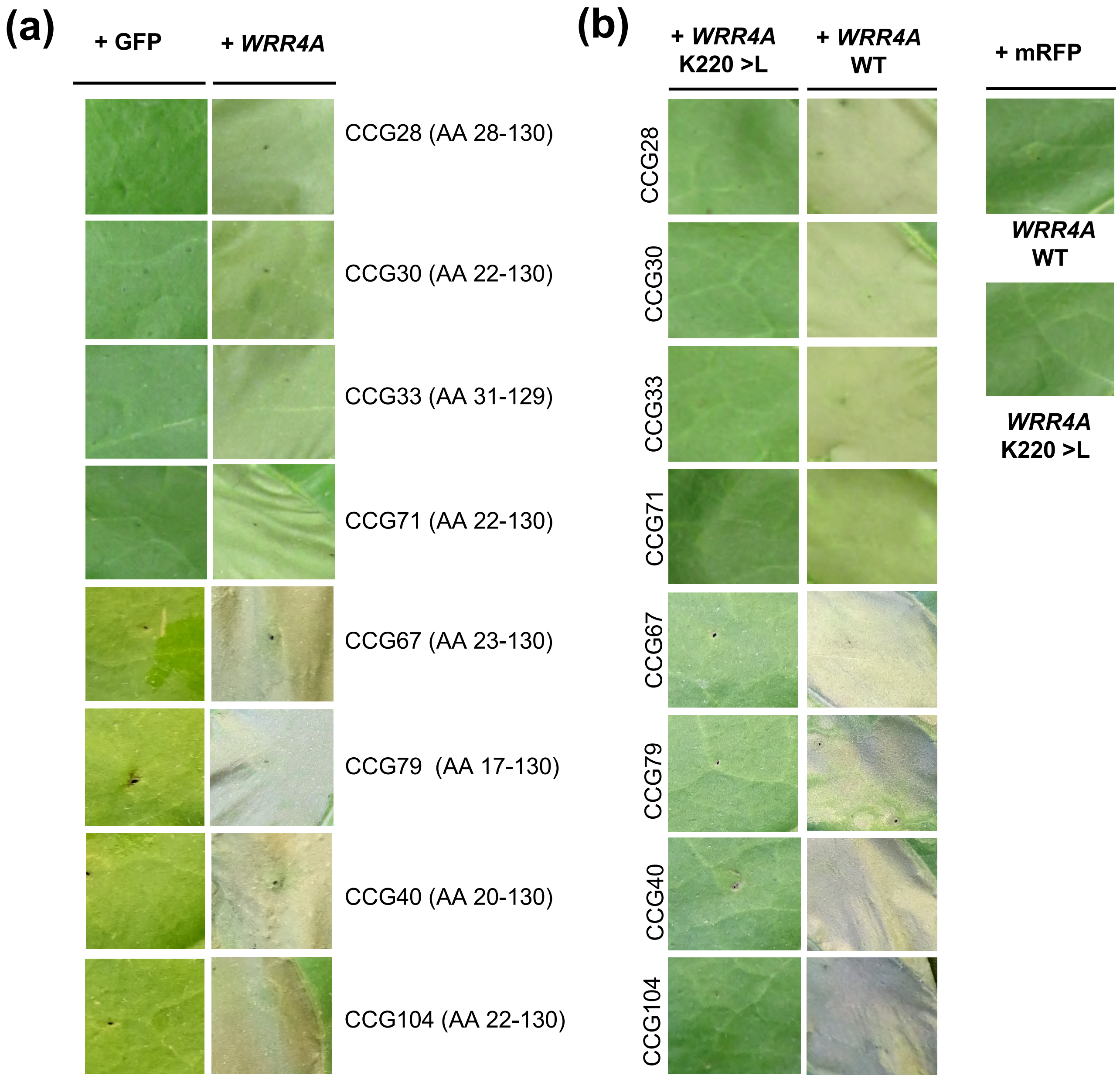


**Fig. S6 CCG N-terminal portion is sufficient for recognition by *WRR4A* and is dependent on intact P-loop motif**

**(a)** Transient expression of different truncated versions of all *WRR4A* recognized CCGs either with GFP or with *WRR4A* in *N. tabacum*. The N-terminal region of all recognized CCGs if sufficient for *WRR4A* recognition.

**(b)** *WRR4A* mediated CCG recognition is dependent on intact Walker A (P-loop) motif.

### Fig. S7

▼ Recognised region of CCG30

```
                               ▼
CCG30_AcNc2     MIHPYWLFLSVVCLRNKVTNAKDPVIITATFSTDSNLGECQFCLTNIAGVNRISLLSITT    60
CCG16_AcNc2     MIHPYWLFLSVVCLRNAVTDAKDPVIITATFSVDSNLGECQFCLTNIAGVDRISLGSIST    60
                **************** **:************.*****************:**** **:*

CCG30_AcNc2     HYSNPPSATAILEAFGLVSSFLRAGIECQGKSKCGSITMTGLEHVNIKPSLETAKKPRCV    120
CCG16_AcNc2     YYSNPPSATAILEAYGWVSSFQRAGIECQGKSKCGFVTMTGLEHVDIDSSLRNMNAPLCV    120
                :*************:* **** ************* :********:*. **.. : * **

                          ▼
CCG30_AcNc2     LTDVQTTFECGVCLQMASGSHMIEKFLLVTDDETKHYFHVLYTTQSNIFEPCSFSGICSG    180
CCG16_AcNc2     LTDVQTTFECGICLQMVSGSHVIEKKQLVTHDRSEHYLHVLYTMQRDIFEPCSFSGICSG    180
                ***********:****.****:***  ***.*.::**:***** * :*************

CCG30_AcNc2     VSYEFADNTCVNLVNKLNQFHFRSVQVAPSSEPKPNLAKMHVAIKAVYDTTLSPSSCFKS    240
CCG16_AcNc2     VNYEFDDSTCVNLINKLTKYHFSSVQVDHLDRPKPKPAKMRVAIKAIDHTTQRTSLCYNT    240
                *.*** *.*****:***.::** ****   ..***: ***:*****: .**   * *:::
```

**Fig. S7 *WRR4A* recognizes N-Terminal portion of CCG30 but not close paralog CCG16**
The role CCG motif is dispensable for recognition by *WRR4A*. The alignment of protein sequences of the two paralogs CCG30 and CCG16 which shows a high identity in their sequence but CCG16 is not recognized by *WRR4A*. The red triangles indicate the minimal recognized region for CCG30.

### Fig. S8

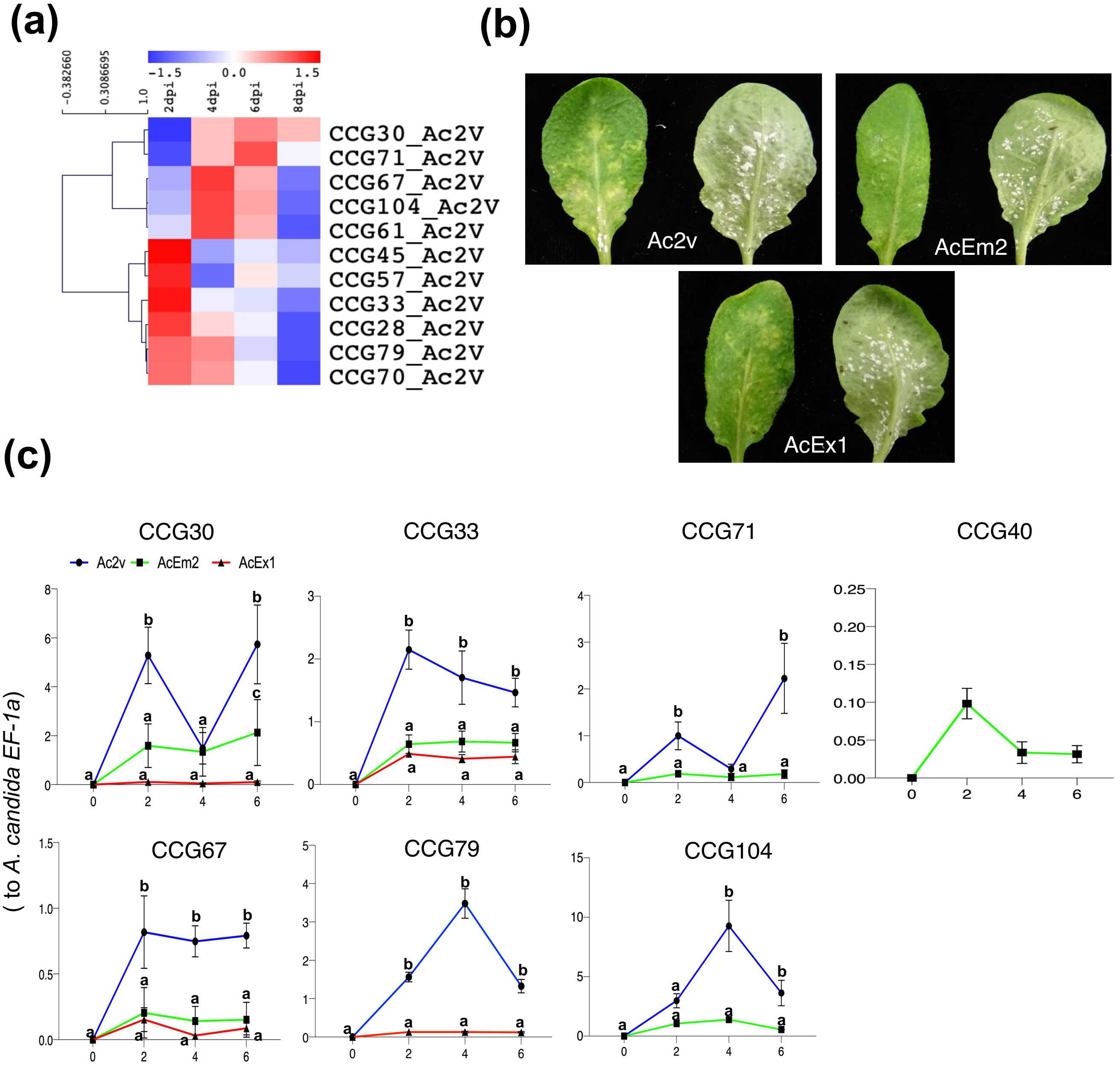


**Fig. S8 Expression profiling of recognized CCGs by RNASeq and RT-qPCR analysis**

(a) Expression profiles of all the *WRR4A*- and *WRR4B*-recognized CCGs from the RNA-Seq data obtained over the consecutive time-points during serial infection stages of *A. candida* Race Ac2V. The recognized CCGs show a clear *in planta* expression suggesting their role during plant colonization.

(b) Disease phenotypes on ws-*eds1 A. thaliana* plants upon infection *A. candida* races Ac2V, AcEm2 and AcEx1.

(c) Transcript levels of the different alleles of *WRR4A* recognized CCGs in *ws-eds1* infected plants at different timepoints after inoculation. Expression was normalized to the Ac*EF1a* gene. Different letters indicate statistically significant differences between the different alleles tested (2 way ANOVA, Bonferroni´s multiple comparison test, $p < 0.05$). Error bars represent SD.

### Fig. S9

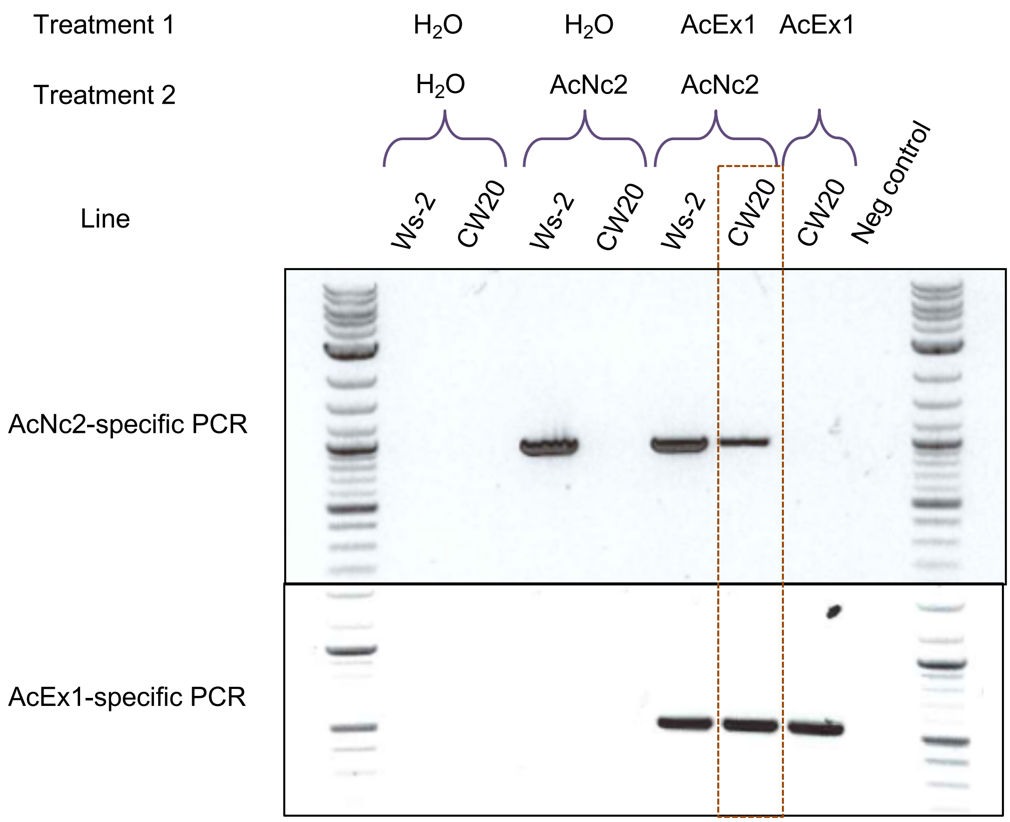


**Fig. S9 AcEx1 can suppress *WRR4A* mediated resistance**
Co-inoculation of AcEx1 and AcNc2 onto CW20 (*WRR4*$^{Col\text{-}0}$) and Ws-2 (*wrr4*) reveals AcEx1 to suppress WRR4 mediated immunity to AcNc2 (observed as AcNc2-specific amplicon present in CW20 pre-inoculated with AcEx1).
