## Supplemental Tables for "The Arabidopsis *WRR4A* and *WRR4B* paralogous NLR proteins both confer recognition of multiple *Albugo candida* effectors": TableS2.pptx

### Slide 1
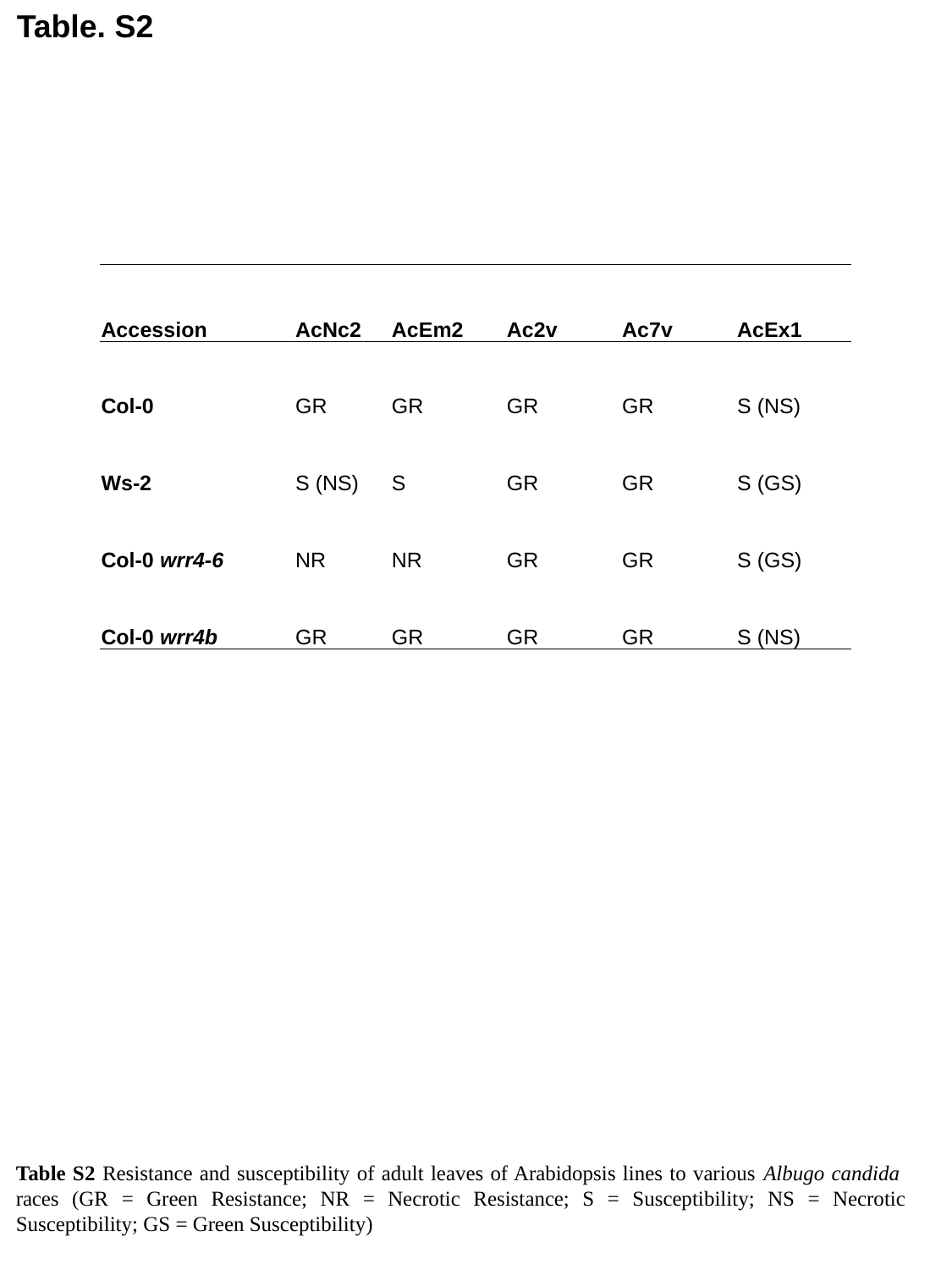

Table. S2
| Accession | AcNc2 | AcEm2 | Ac2v | Ac7v | AcEx1 |
| --- | --- | --- | --- | --- | --- |
| Col-0 | GR | GR | GR | GR | S (NS) |
| Ws-2 | S (NS) | S | GR | GR | S (GS) |
| Col-0 wrr4-6 | NR | NR | GR | GR | S (GS) |
| Col-0 wrr4b | GR | GR | GR | GR | S (NS) |
Table S2 Resistance and susceptibility of adult leaves of Arabidopsis lines to various Albugo candida races (GR = Green Resistance; NR = Necrotic Resistance; S = Susceptibility; NS = Necrotic Susceptibility; GS = Green Susceptibility)
